# Kinesin-12 KLP-18 contributes to the kinetochore-microtubule poleward flux during the metaphase of *C. elegans* one-cell embryo

**DOI:** 10.1101/2022.11.07.515476

**Authors:** Nina Soler, Mathis Da Silva, Christophe Tascon, Laurent Chesneau, Pauline Foliard, Hélène Bouvrais, Sylvain Pastezeur, Loïc Le Marrec, Jacques Pecreaux

## Abstract

The mitotic spindle, a key structure to partition chromosomes during cell division, connects its poles to the chromosomes through microtubules. Their plus-ends, oriented towards the chromosomes, exhibit dynamic instability crucial for kinetochore correct attachments. Involved in this process, the poleward flux implicates the displacement of microtubules towards the spindle poles, coordinated with polymerisation at the plus ends. The mechanisms behind this are diverse. It includes treadmilling powered by microtubule depolymerisation at the spindle poles, sliding of spindle microtubules by molecular motors like Kinesin-5, and pushing microtubules away from the chromosomes by chromokinesins. Interestingly, no such flux was reported in the *Caenorhabditis elegans* zygote, although all proteins contributing to flux in mammals have homologs in the nematode.

To explore this, we fluorescently labelled microtubules and conducted photobleaching. We found no global poleward flux; the bleached zone’s edges moved inward. The centrosome-side front motion was caused by dynamic instability, while the chromosome-side front exhibited faster recovery, suggesting an additional mechanism. This larger slope was detected only near the chromosomes, indicating that only kinetochore microtubules undergo flux. Consistently, this flux depended on proteins ensuring the chromosome attachment and growth of the kinetochore microtubules, notably NDC-80, CLS-2^CLASP^, and ZYG-9^XMAP215^. Furthermore, this flux decreased as metaphase progressed and attachments transitioned from side-to end-on; it was reduced by SKA-1 recruitment. Treadmilling was unlikely to account for these observations, as most kinetochore microtubules do not reach spindle poles in the zygote spindle. Conversely, the depletion of kinesin-12 KLP-18^KIF15^, which cross-links and focuses microtubules at meiosis, reduced the front rate. Ultimately, we propose that the sole kinetochore microtubules slide along spindle microtubules, likely powered by KLP-18, contrasting with solid displacement in other systems. It aligns with observations in human cells of decreasing flux with increasing chromosome distance.

## INTRODUCTION

During cell division, the mitotic spindle ensures a faithful partitioning of the genetic material into two identical sets distributed to each daughter cell. This specialised structure is assembled from microtubules (MTs) and their associated proteins (MAPs), especially molecular motors and cross-linkers. The microtubules are polarised semi-flexible polymers which alternate growth and shrinkage, and their plus ends are more dynamic than their minus ends (Akhmanova and Steinmetz, 2015). In the nematode C. *elegans* zygote, there are two types of microtubules in the spindle: kinetochore microtubules (kMT) with plus-ends anchored at the chromosomes and minus-ends spread along the spindle; and spindle microtubules (sMT) with minus-ends at the centrosomes and plus-ends not contacting the kinetochore (Muller-Reichert et al., 2010; Redemann et al., 2017). We included the branched microtubules along the sMT lattice in this latter category. Microtubule dynamic instability is essential for the spindle functions, a feature largely used in cancer therapies (Steinmetz and Prota, 2018; Vicente and Wordeman, 2019; Wordeman and Vicente, 2021). Numerous MAPs ensure a precise regulation of microtubule dynamics along the various phases of mitosis and spindle zones (Lacroix et al., 2018; Roostalu et al., 2018; Srayko et al., 2005). These molecular motors and MAPs stochastically bind and unbind to the microtubules, making the spindle highly dynamic (Elting et al., 2018; Nazockdast and Redemann, 2020).

An important aspect of ensuring spindle function is the attachment of the chromosomes by the kMTs while preserving the dynamic instability of the microtubule plus-ends (Bakhoum and Compton, 2012). Thus, a specialised structure assembles at the centromeric regions of the chromosomes: the kinetochore (Cheeseman, 2014; Musacchio and Desai, 2017). In particular, the outer kinetochore protein NDC-80 makes the bridge with the microtubules (Cheeseman et al., 2004; Suzuki et al., 2016; Ye et al., 2016). In the nematode, chromosomes are holocentric; kinetochores are scattered all along the chromosomes (Maddox et al., 2004). Chromosome attachment first occurs laterally, along the microtubule lattice (side-on), before being converted into an attachment at the plus-end (end-on) (Cheerambathur et al., 2013; Magidson et al., 2011). This latter can withstand the tension from the microtubules on the kinetochore. This tension is essential for the spindle assembly checkpoint (SAC) to identify chromosome misattachment and also for the subsequent mechanisms to fix these mistakes (Kuhn and Dumont, 2019; McVey et al., 2021). When attachments are correct, they are secured by the SKA complex, and the kMT dynamicity is reduced (Cheerambathur et al., 2013; Cheerambathur et al., 2017; Hanisch et al., 2006; Schmidt et al., 2012). MAPs regulate the dynamics of the attached microtubule, like the rescue factor CLS-2^CLASP^ (Al-Bassam and Chang, 2011; Cheeseman et al., 2005; Matthews et al., 1998). Some ubiquitous regulators also contribute, such as the polymerisation enhancer ZYG-9^XMAP215^ (Matthews et al., 1998). In the nematode, like in other organisms, only one-fifth of the kMTs can directly reach the spindle pole (Kiewisz et al., 2022; Petry, 2016; Redemann et al., 2017). The remaining kMTs are likely connected to sMTs since their minus ends are very close to the sMT lattices.

Capturing and attaching chromosomes are partly stochastic, which can lead to errors. These are usually intercepted by the SAC (Nicklas, 1997). Notably, as in mammalian embryos, the C. *elegans* zygote has a weak SAC, and some of its components are dispensable (Gerhold et al., 2018; Oegema and Hyman, 2006; Pintard and Bowerman, 2019; Tarailo et al., 2007). It, however, relies on the tension exerted by the microtubules on the kinetochore, which in turn involves their polymerisation (Bakhoum et al., 2009; Cimini et al., 2006; Ertych et al., 2014; Lampson and Grishchuk, 2017; Schwietert et al., 2022). Combined with minus-end depolymerisation, it leads to poleward flux, which is a displacement of the microtubule network away from chromosomes (Barisic et al., 2021; Hotani and Horio, 1988; Mitchison, 1989; Steblyanko et al., 2020; Waters et al., 1996). A poleward flux was measured in various organisms, from Xenopus to mammals (Brust-Mascher et al., 2004; Brust-Mascher et al., 2009; Maddox et al., 2002; Maddox et al., 2003; Maiato et al., 2005; Matos et al., 2009; Yang et al., 2008). It plays an important role in correcting chromosome attachment errors, putatively by renewing the microtubules in contact with the kinetochore (Ertych et al., 2014; Ganem et al., 2005; Matos et al., 2009; Pereira and Maiato, 2012). During anaphase A, it also contributes to a synchronous migration of the sister chromatids toward their facing spindle poles (Ganem et al., 2005; Matos et al., 2009; Mitchison and Salmon, 1992). The most obvious flux mechanism is treadmilling, in which the microtubule grows on the kinetochore side in a coordinated fashion with shortening at the spindle pole; this latter process powers the mechanism (Gaetz and Kapoor, 2004; Ganem et al., 2005). Such a mechanism was also recreated *in vitro* following a minimal system approach with only four proteins: microtubule polymerase XMAP215^ZYG-9^, the plus-end binding protein EB1^EBP-2^, the rescue factor CLASP2^CLS-2^, and the microtubule depolymerising kinesin MCAK^KLP-7^ (Arpag et al., 2020). This latter kinesin is essential to depolymerise the microtubules on the centrosome side. It can be assisted by severing enzymes like katanin, spastin and fidgetin (Zhang et al., 2007). Furthermore, treadmilling is consistent with the gel-like behaviour of the spindle reported in Xenopus egg extracts (Dalton et al., 2022) and the classic flux measurements using photobleaching/photoconversion experiments (Barisic et al., 2021; Mitchison and Salmon, 1992). Their results suggest a mostly solid motion of the spindle microtubule network at the steady state. Beyond the treadmilling mechanism, alternatives are twofold. Firstly, specialised molecular motors like kinesin-5 KIF11^BMK-1^ (also known as EG-5) and kinesin-12 KIF15^KLP-18^ can slide the overlapping antiparallel microtubules carrying the kMTs through cross-linking (Miyamoto et al., 2004; Steblyanko et al., 2020; Uteng et al., 2008). However, kinesin-12 seems more specialised in parallel microtubules (Drechsler and McAinsh, 2016). Variants were proposed by sliding bridging fibres along each other (Jagric et al., 2021). The motion of these fibres is transmitted to other microtubules, including the kMTs and the sMTs, by cross-linking agents or motors like HSET^KLP-15/16/17^, NuMA^LIN-5^ or PRC1^SPD-1^ (Elting et al., 2014; Risteski et al., 2022; Steblyanko et al., 2020; Wang et al., 2025). Secondly, chromokinesin KIF4A^KLP-19^ congresses the chromosome arms and generates a reaction force that pushes the kMTs away from the chromosomes (Steblyanko et al., 2020; Wandke et al., 2012). It may play a similar role in nematodes, although it would also regulate microtubule dynamics (Powers et al., 2004; Zimyanin et al., 2023). These various mechanisms can also be superimposed to ensure a robust poleward movement of the kMTs.

Surprisingly, although homologous proteins are present in the nematode C. e*legans,* no flux was reported in anaphase, while a putative but weak anti-poleward flux was suggested at metaphase (Labbe et al., 2004; Redemann et al., 2017). Notably, microtubules are highly dynamic in the nematode due to tubulin specificities (Chaaban et al., 2018). In contrast, a flux was found in zygote meiosis (Lantzsch et al., 2021). These paradoxes led us to investigate the flux during mitotic metaphase using the established fluorescence recovery after photobleaching (FRAP) (Axelrod et al., 1976; Giakoumakis et al., 2017; Matsuda and Nagai, 2014; White and Stelzer, 1999). Whereas we found no net flux for the spindle as a whole, we measured a poleward displacement of the microtubules close to the chromosomes. It depends on kinetochore proteins like CLS-2^CLASP^, NDC-80 and the kinetochore-attachment states. We finally suggest that the kMTs be slid along the fixed sMTs by KLP-18^KIF15^.

## RESULTS

### Two distinct microtubule-dynamics mechanisms account for recovering fluorescence after photobleaching

We investigated the dynamics of microtubules within the metaphasic spindle of the *C. elegans* zygote, studying recovery after photobleaching a 2.6µm-wide region in the spindle (Methods). We used the GFP::TBB-2^β-tubulin^ labelling. We previously reported no phenotype for this strain (Bouvrais et al., 2021). By analysing the average intensity in the photobleached region over time, we excluded that the diffusion of tubulin dimers accounted for the recovery inside the spindle (Supplemental Text §1 and Fig 1AB). Next, to investigate a putative flux, we set out to analyse the shape of the bleached area over time (Movie S1, Fig 1CD). We used the centrosome on the bleaching side as a spatial reference to register images and to remove the effects of spindle pole motion (Methods § Image processing, Fig S2). We produced the kymograph of individual embryos along the spindle axis by median projection along the transverse axis (Fig S3). The fast dynamics of the microtubules necessitated imaging at 12.5 frames per second, resulting in a low signal-to-noise ratio (SNR). To account for low SNR, we averaged the kymographs from several embryos after aligning them on the centrosome-side top corner of the bleached region and matching their intensity histograms. We then segmented the bleached region of the averaged kymograph. We fitted the boundaries of the segmented region with a line to quantify the displacement of the two fronts over time by their slope in the kymograph (Fig. 1F, blue lines). We used a leave-one-out resampling (Jackknife) to obtain the standard errors of the slopes (Efron and Tibshirani, 1993; Quenouille, 1956). Such an approach also enabled us to visualise the front-slope distributions by displaying N-1 averages (Fig 1E, Methods § Statistics on kymograph front slopes). We finally computed the slope of the mid-curve between the edges corresponding to the bleached region solid displacement (Fig 1F, black line). It read 0.015 ± 0.023 µm/s, oriented poleward (*N =* 10, mean ± standard error (se) obtained by Jackknife resampling, *p* = 0.68 compared to 0). We concluded that the bleached area underwent no global displacement, meaning no significant global flux. This is consistent with previous reports on the lack of flux in metaphase and anaphase (Labbe et al., 2004; Redemann et al., 2017).

**Figure 1:**
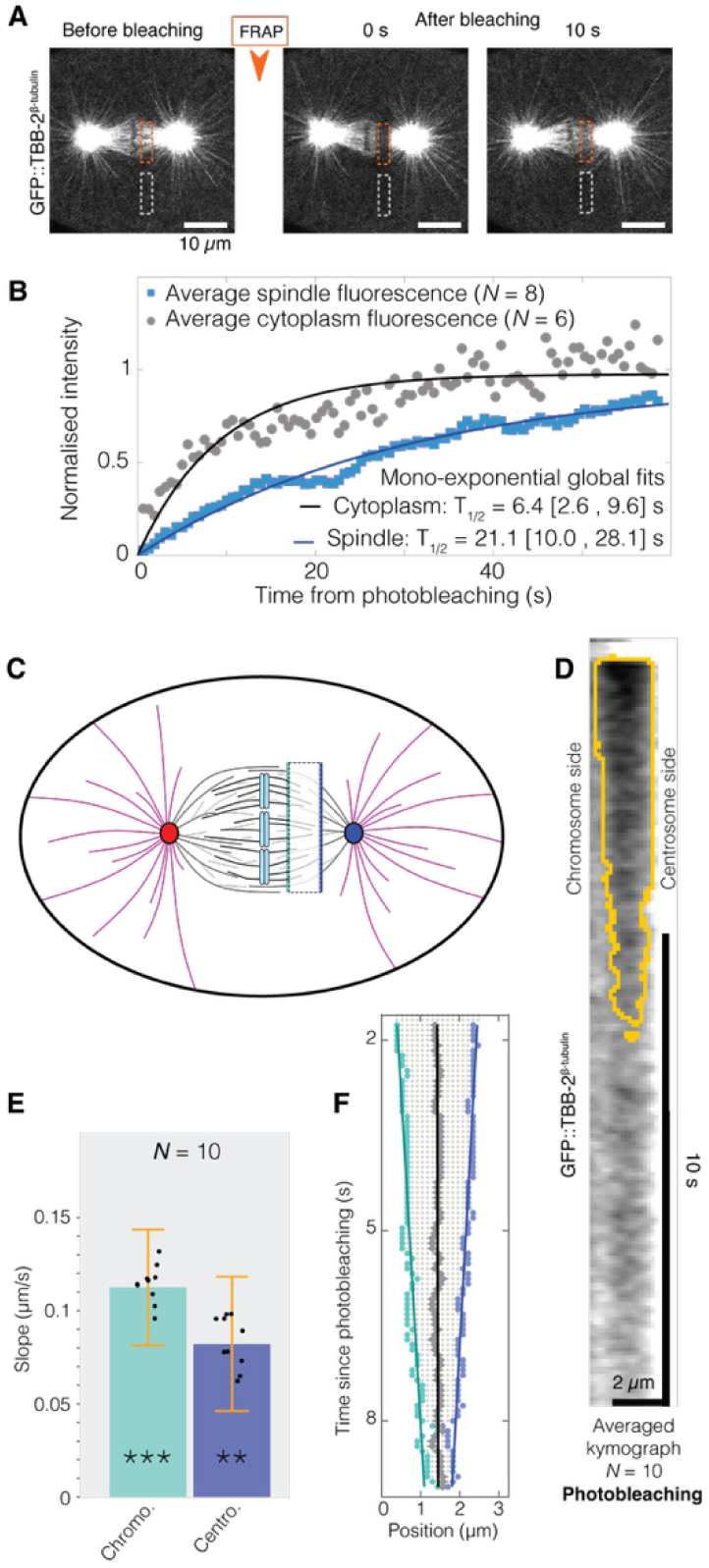
Tubulin fluorescence recovery after photobleaching in the metaphasic spindle of *C. elegans*. (**A**) Representative confocal live images before and after photobleaching (time above the stills). The posterior centrosome is on the right-hand side. Microtubules are labelled with GFP::TBB-2^β-tubulin^. The scale bar corresponds to 10 µm. Orange and grey dashed boxes depict the regions used to assess fluorescence recovery in the spindle and cytoplasm, respectively. (**B**) Fluorescence recovery averaged over GFP::TBB-2^β-tubulin^ non-treated embryos bleached: (blue) *N* = 8 in the spindle and (grey) *N* = 6 in the cytoplasm. Lines correspond to the global fit by a single exponential model (Giakoumakis et al., 2017) (Suppl Text §1). The corresponding half-lives are reported with 95% confidence intervals in brackets. (**C**) Schematics of the FRAP experiment to measure front displacements. Thick dark grey lines depict the spindle microtubules emanating from the (red) anterior and (blue) posterior centrosomes; thin light grey lines depict the one branching from other microtubule lattices. Black lines correspond to kinetochore microtubules bound to (blue bars) the condensed sister chromatids. Astral microtubules are depicted in purple colour. The dashed box corresponds to the bleached area. The blue bars represent the measurement locations of the fronts on (dark blue) the chromosome and (light blue) the kinetochore sides. (**D**) Averaged kymograph over the posterior half-spindle of *N =* 10 GFP::TBB-2^β-tubulin^ labelled embryos. The centrosome was located on the right-hand side. The orange line delineates the bleached region as obtained by our analysis (Movie S1, Methods § Image processing). (**E**) Front velocities by analysing the segmented bleached-region of *N =* 10 GFP::TBB-2^β-tubulin^ non-treated embryos. Black dots represent the average of *N*-1 embryos, leaving out each embryo in turn (Methods § Statistics on kymograph front slopes). Overlaid stars indicate significance relative to 0 obtained by jackknife resampling (Methods § Statistics). Bars correspond to means; error bars are estimated standard errors using Jackknife resampling. Light blue bars represent velocities on the chromosome side, and dark blue bars represent those on the centrosome side. It corresponds to the kymograph analysis displayed in B. (**F**) Linear fits of the edges of the segmented kymograph averaged over the non-treated embryos reported in panels B and C. Grey dots depict the pixels in the bleached region. Boundaries are highlighted by coloured dots on (turquoise, left edge) the chromosome side and (blue, right edge) the centrosome side. Grey diamonds depict the region mid-line. Linear fits are reported as lines of corresponding colours.

However, a visual inspection of the kymograph (Fig 1E, F) indicated a fluorescence recovery by closing the edges through two inward-oriented fronts. We investigated this V-shaped recovery and measured the closure velocities, both oriented inwards (Fig 1E). Chromosome and centrosome side values were significantly larger than 0. We reckoned that these two fronts, moving in opposite directions, might reveal distinct microtubule dynamics close to and far from the chromosomes. Indeed, we expect the kMTs to be numerous enough to be detected near chromosomes but to be vastly overwhelmed by sMTs at the position where the centrosome front is measured (Redemann et al., 2017).

To safeguard against artefacts, we repeated the experiment using photoconversion (see Methods § Microscopy). We crossed strains mEOS3.2::TBB-2^β-tubulin^, which labels microtubules, and mCherry::TBG-1^γ-tubulin^, which reveals centrosomes. We used this second label to register the images over time, as reported above (Methods § Image processing). We produced the kymographs, and again, to cope with the low SNR, we averaged them across embryos after aligning them on the photoconverted region. We obtained a similar V-shaped pattern (Fig S1A) as expected. We concluded that the two fronts were not artefactual.

We set out to validate that the motion of the two closing edges depended on microtubule dynamics. We used a mild *zyg-9*^XMAP215^*(RNAi)* to preserve an apparently normal spindle, although leading to a shorter spindle: 12.16 ± 0.22 µm (*N* = 11, t-test *p* = 1.4×10^-5^) compared to 13.71 ± 0.17 (*N* =13) in control RNAi embryos. We investigated the front on the centrosome side (Suppl Text §2) (Srayko et al., 2003). This depletion decreases the polymerisation rate at the plus ends (Srayko et al., 2005). We observed a 13% and 20% reduction in slope on the chromosome and centrosome sides, respectively, though neither was significant (Fig. S1BC, replicated in Fig. S1D).

The most obvious way to account for the observed fronts is through a flux of microtubules themselves. While a poleward flux of kMTs was not observed in the nematode, it is well-known in numerous other species. In contrast, an anti-poleward flux of sMTs appeared much less likely. The edge of the bleached region on the centrosome side was located at about half the distance between the chromosomes and the spindle pole; at this place, the sMTs are three times more numerous than the kMTs (Redemann et al., 2017). We thus focused on the sMTs and reasoned that the front displacement on the centrosome side might be accounted for by the dynamics of the sMTs (growth and shrinkage) combined with diffraction from imaging. We modelled this phenomenon and computed a closing front similar to the experimental one (Suppl Text §3, Fig 2A, S5). In particular, our model predicted a reduction in the centrosome-side front slope as the spindle got longer (Fig2B). Intuitively, it takes more time for the sMTs to regrow from the spindle poles in a longer spindle, since the pole-to-bleached-area distance is larger. We observed experimentally such a dependence specifically on the centrosome side (Fig 2C, Suppl Fig S4), by considering non-treated and all control RNAi from the present study, taking advantage of the natural length variability (Le Cunff et al., 2024). Interestingly, we did not observe such a correlation on the chromosome side, and we attributed that to an additional mechanism superimposed on that side. We concluded that dynamic instability is sufficient to account for the front motion on the centrosome side. However, because no clear correlation was observed between the chromosome side slope and the spindle length (Fig S4), and because recovery was faster on that side, it suggested an additional mechanism on the chromosome side.

**Figure 2:**
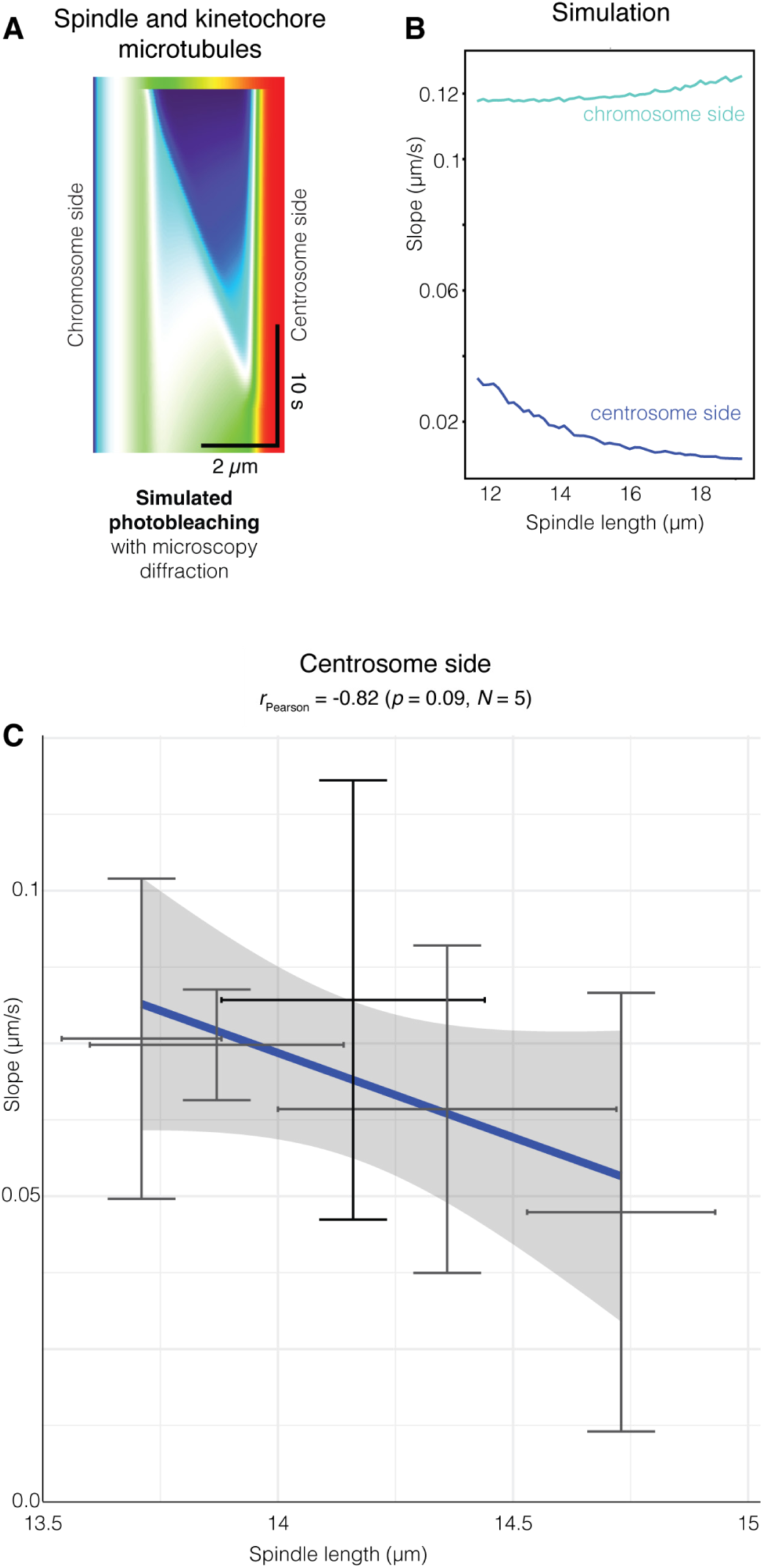
Front velocity on the centrosome side did not require flux but depended on spindle length. (**A**) Simulated kymograph including not-fluxing sMTs, and kMTs submitted to poleward flux (Suppl Text §3). It accounts for diffraction by microscopy imaging. Corresponding simulated slopes read 0.12 µm/s on the chromosome side and 0.016 µm/s on the centrosome side. The colours scale ranges from blue for dark pixels to red for bright areas. Simulation parameters are reported in Suppl Table S1. (**B**) The model predicted an anti-correlation between (blue) the centrosome-side front slope and the spindle length. (turquoise) In contrast, no correlation was predicted on the chromosome side, matching the experimental observation. **(C)** The experimental front velocities on the centrosome side in (black line) non-treated and (grey line) RNAi control conditions did not show a significant correlation with the spindle length. We used strains with labelled microtubules GFP::TBB-2^β-tubulin^. Spindle length was measured at the time of bleaching. Pearson correlation coefficient and the corresponding test are indicated above the plot. Bars correspond to means; error bars are standard errors. The grey-shaded region corresponds to the 95% confidence interval on the line coefficients.

### A poleward flux of kinetochore microtubules may account for the chromosome-side recovery

We reckoned that the higher front slope observed on the chromosome side and the lack of correlation between the spindle length and the recovery front slope suggested that a second mechanism might superimpose specifically on the chromosome side. The most obvious addition would involve the kMTs growing at their plus ends, attached to the chromosomes, while sliding towards the poles and thus along the spindle microtubules. Consistently, depletion of ZYG-9^XMAP215^ reduced the front slope on the chromosome side (Fig S1B, D). This hypothesis was consistent with Redemann and colleagues not measuring this poleward flux because they were far from the chromosomes. Indeed, the kMTs are abundant compared to the sMTs, only close to the chromosomes (Redemann et al., 2017).

In other organisms, the poleward flux is reported to involve all microtubules, although their dynamics may differ (Barisic et al., 2021). Dalton and co-authors proposed that the spindle undergoes gelation to account for the solid poleward motion of the microtubules within spindles prepared from *Xenopus laevis* extracts (Dalton et al., 2022). In contrast, we proposed that only the kMTs would undergo a poleward flux. As their number decreases with distance from the kinetochore (Redemann et al., 2017), the flux could be seen as restricted close to the kinetochores. To challenge this idea, we performed FRAP experiments, bleaching a smaller area either close to or far from the spindle pole (Fig 3A). When looking far from the chromosomes, we obtained similar front velocities on edges facing chromosomes and centrosome (Fig 3B, maroon shading). It suggested that, in this spindle region, both fronts reflected similar mechanisms. Moreover, this symmetry is incompatible with sMTs undergoing a flux in either direction. Because the sMTs are in the vast majority compared to the kMTs, the recovery was likely due solely to the microtubule dynamics combined with imaging diffraction, as detailed above. Close to the kinetochore, we measured a significantly steeper front slope on the chromosome side than on the centrosome side, by 15%, consistent with our previous measurement (Fig 3B, grey shading). However, the slope on the chromosome side was larger than in the bleached region far from the centrosomes. It may be related to a gradient of RAN protein around chromosomes that promotes microtubule growth (Bamba et al., 2002; Srayko et al., 2005). Overall, this experiment was consistent with our hypothesis that only the kMTs underwent poleward flux. It also accounted for the difference between our findings and previous measurements, as the kMT flux could be detected only close to the chromosomes (Labbe et al., 2004; Redemann et al., 2017). Anywhere else, the non-fluxing sMTs, which are in the majority, hide the kMTs flux.

**Figure 3:**
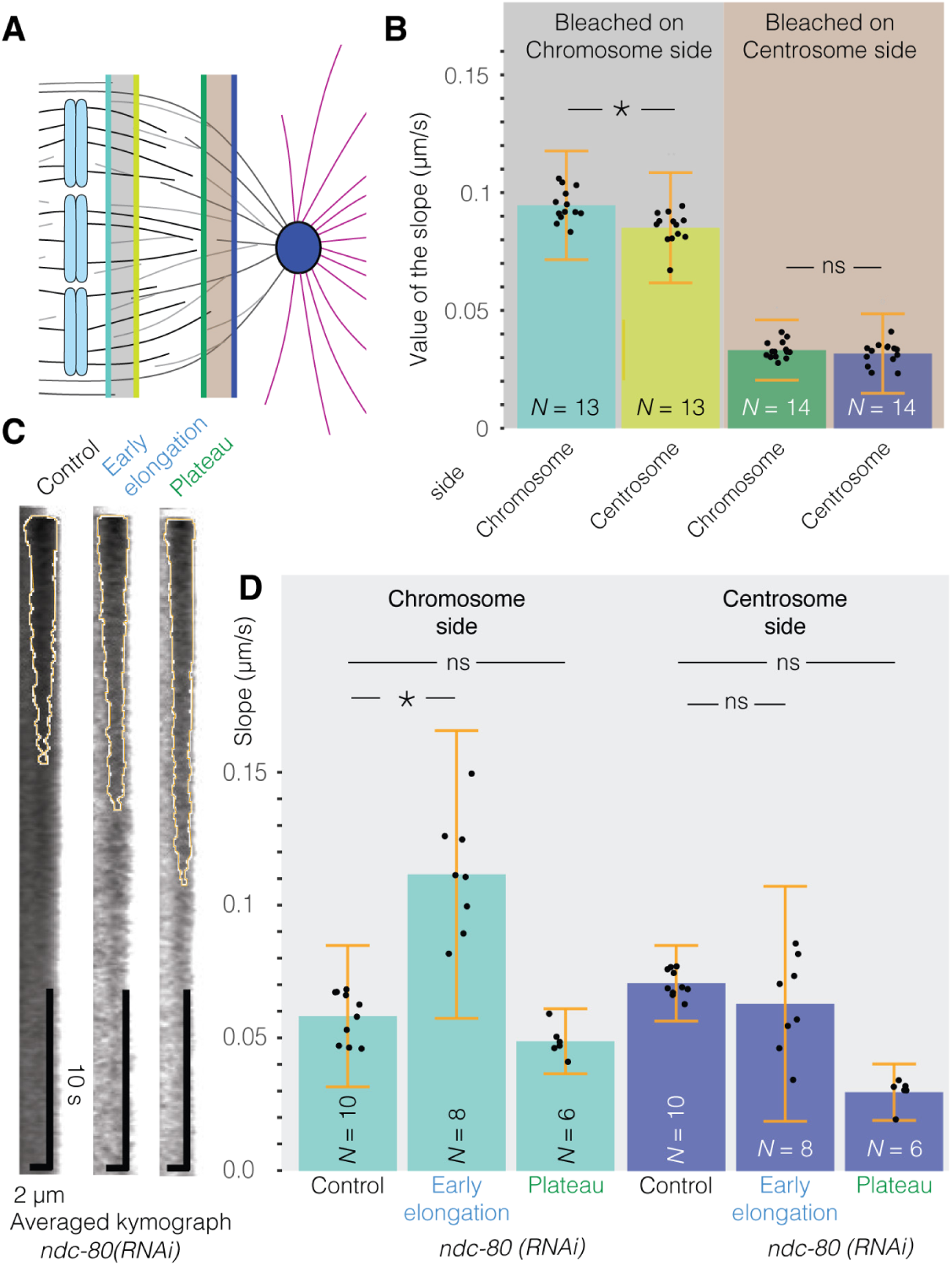
Front velocity on the chromosome side requires kMT polymerising at the plus-ends. (**A**) Schematics of the FRAP experiment, with bleaching localised close to either the chromosomes or the centrosome to measure front displacement during recovery. Grey lines depict the spindle microtubules emanating from (blue) the posterior centrosome. Black lines correspond to the kinetochore microtubules bound to (blue bars) the condensed sister chromatids. The grey and maroon boxes mark the bleached areas near the chromosomes and the centrosome, respectively. The coloured lines depict the approximate positions of the front measurements corresponding to B. (**B**) Front velocities by segmenting the dark regions of non-treated embryos, bleached either (grey background, *N* = 13) close to chromosomes or (maroon background, *N* = 14) close to centrosomes. The colour of the bars corresponds to the schematics (A). (**C**) Averaged kymographs over the posterior half-spindle for the data presented in (D), with centrosome on the right-hand side. The orange line delineates the bleached region as obtained by our analysis (Methods § Image processing). (**D**) Front velocities by segmenting the dark region of *N =* 8 *ndc-80(RNAi)* embryos bleached during early elongation, *N =* 6 *ndc-80(RNAi)* bleached during the spindle length plateau and their *N* = 10 controls (typical micrograph at Fig S6A). Light blue bars are values on the chromosome side and dark blue bars on the centrosome side. The experiment was replicated (Fig S6C). In all cases, microtubules were labelled using GFP::TBB-2^β-tubulin^. In panels B and D, black dots represent averages of *N*-1 embryos, leaving out, in turn, each embryo (Methods § Statistics on kymograph front slopes). Bars correspond to means, and error bars are estimated using Jackknife resampling.

To further ascertain that fluxing kMTs could cause the front motion on the chromosome side, we set out to genetically impair the kMTs’ attachment to the kinetochore. We partially depleted NDC-80 under conditions that allowed chromosome segregation eventually to succeed in anaphase (Methods). We observed that the spindle in metaphase elongated during early metaphase (further termed early elongation) and plateaued before the anaphase onset, as previously reported (Fig S6A) (Cheerambathur et al., 2013). We studied the microtubule dynamics by FRAP during both phases (Fig 3C, D). The control embryos were imaged throughout the last two minutes of metaphase, as the normal spindle’s slight metaphase elongation does not exhibit two phases. Interestingly, only the chromosome-side front was significantly increased, by 92% during early elongation, suggesting that the kinetochore-microtubule attachment was involved in setting the corresponding front slope. We repeated the experiment and found an increased slope again during early elongation, by 45%, confirming our hypothesis (Fig S6C). We reasoned that our partial depletion of NDC-80 led to a delay in microtubules attaching end-on to kinetochores (Cheerambathur et al., 2013; Cheerambathur et al., 2017; Cheeseman et al., 2004; Lange et al., 2019). Thus, kinetochore-microtubule attachments could not withstand normal load, leading to microtubule snatch-out, as evidenced by early elongation, under the hypothesis that the kMTs underwent a poleward flux. Later, during the plateau, we observed a slope lower than that of the control. Altogether, these observations were suggestive that growing kMTs at the kinetochores may be necessary for the mechanism causing the chromosome recovery front.

To further relate microtubule plus-end dynamics to the measured front slope, we reasoned that, in the classic poleward-flux mechanism, CLS-2^CLASP^ is critical to ensure that microtubules switch to polymerisation during flux (Arpag et al., 2020; Cheeseman et al., 2005). We thus depleted this protein, but only partially, to prevent spindle breakage. We observed a shorter spindle during metaphase, as expected (Cheeseman et al., 2005) (Fig S6B, F). We monitored the front slope using our FRAP assay and found a slope increased by 23% (170% in replica) on the chromosome side that recalled the one observed during the precocious spindle elongation upon *ndc-80(RNAi)* (S6D, replicated in S6E). We interpreted it similarly: the kMTs were pulled off the kinetochores upon *cls-2(RNAi)*, resulting in an increased front slope, although penetrance variability led to phenotypic variability. Consistently, upon strong depletion, the spindle breaks due to the cortical pulling forces (Cheerambathur et al., 2017; Cheeseman et al., 2005). Targeting CLS-2 disturbed the metaphasic plate, leading to somewhat inaccurate slope measurements. Overall, we concluded that our results, especially the observed recovery front on the chromosome side, could be explained by the hypothesis that kinetochore-microtubule fluxes towards the poles, requiring microtubule polymerisation at the plus-ends.

We reasoned that under such a hypothesis, kMTs minus ends should displace towards the poles. We thus imaged the mitotic spindle and focused on the microtubule minus-ends using GFP::ASPM-1 (Connolly et al., 2015; Ellefson and McNally, 2011; Harvey et al., 2023; Jiang et al., 2017; Mullen and Wignall, 2017; Taylor et al., 2023) (Suppl Text §5). Consistently, C. *elegans* ASPM-1 was seen concentrated at the spindle poles in meiosis. We imaged this strain using a confocal microscope with deconvolution (Methods § Microscopy, Movie S3, Fig. S7A-C). Because the ASPM-1 spots were too faint to be tracked, we analysed the optical flow in the mitotic spindle halves with the approach published by Drechsler and colleagues, considering the last 30 s of the metaphase (Drechsler et al., 2020) (Suppl Fig S7D). ASPM-1::GFP displayed a significant inwards (centripetal) optical flow around both centrosomes, consistent with our hypothesis of the kMTs sliding poleward along the sMTs (Fig S7EF, statistics in Suppl Text §5). As a positive control, we imaged EBP-2::mKate2, which forms comets at the plus ends of growing microtubules (Srayko et al., 2005), and observed a significant outwards (centrifugal) optical flow (Suppl Fig S7GH). Tubulin optical flow showed no clear directionality, offering a negative control (Suppl Fig S7IJ). We concluded that a fraction of microtubule minus ends, attributable to a subset of kMTs, was moving towards the spindle poles, supporting our hypothesis.

### The chromosome-side poleward flux depends on the attachment status at the kinetochore

We reckoned that during spindle assembly, kinetochores first attached laterally, with reduced resistance to tension, then end-on (Cheerambathur and Desai, 2014). This second configuration couples the dynamics of the kMT plus-ends to tension, reducing the growth rate (Cheeseman et al., 2004; Suzuki et al., 2015). Next, the SKA complex is recruited during late metaphase, further reducing the dynamicity of the attachments (Cheerambathur et al., 2017; Schmidt et al., 2012). To investigate such a dependence in our measurements, we grouped our non-treated and control experiments into 15s-long bins based on the time the bleaching was performed and measured the front velocities. On the chromosome side, we found that it decreased significantly over time (Fig 4A). It suggested that the dynamics of the kMT plus-ends might impact the measured velocity of the chromosome-side front, which aligned with our hypothesis. We excluded that the observed correlation could result from modest variations in the distance between the bleached region and the chromosomes (Fig S8B). Notably, the front of the centrosome side also displayed a decreasing velocity with metaphase progression (Fig S8A). In the framework of the proposed model for this side (Suppl Text §3), we attributed this to either the decreasing microtubule growth rate over metaphase and anaphase, or the spindle getting slightly longer over metaphase (Goshima and Scholey, 2010; Srayko et al., 2005). Interestingly, we observed decreases in front velocities by 20% and 18% on the chromosome and centrosome sides, respectively, during early anaphase compared to metaphase (Fig S9A). It is consistent with locking correct chromosome attachment at the kinetochores and the reduced microtubule growth rate at anaphase (Asbury, 2017; Srayko et al., 2005).

**Figure 4:**
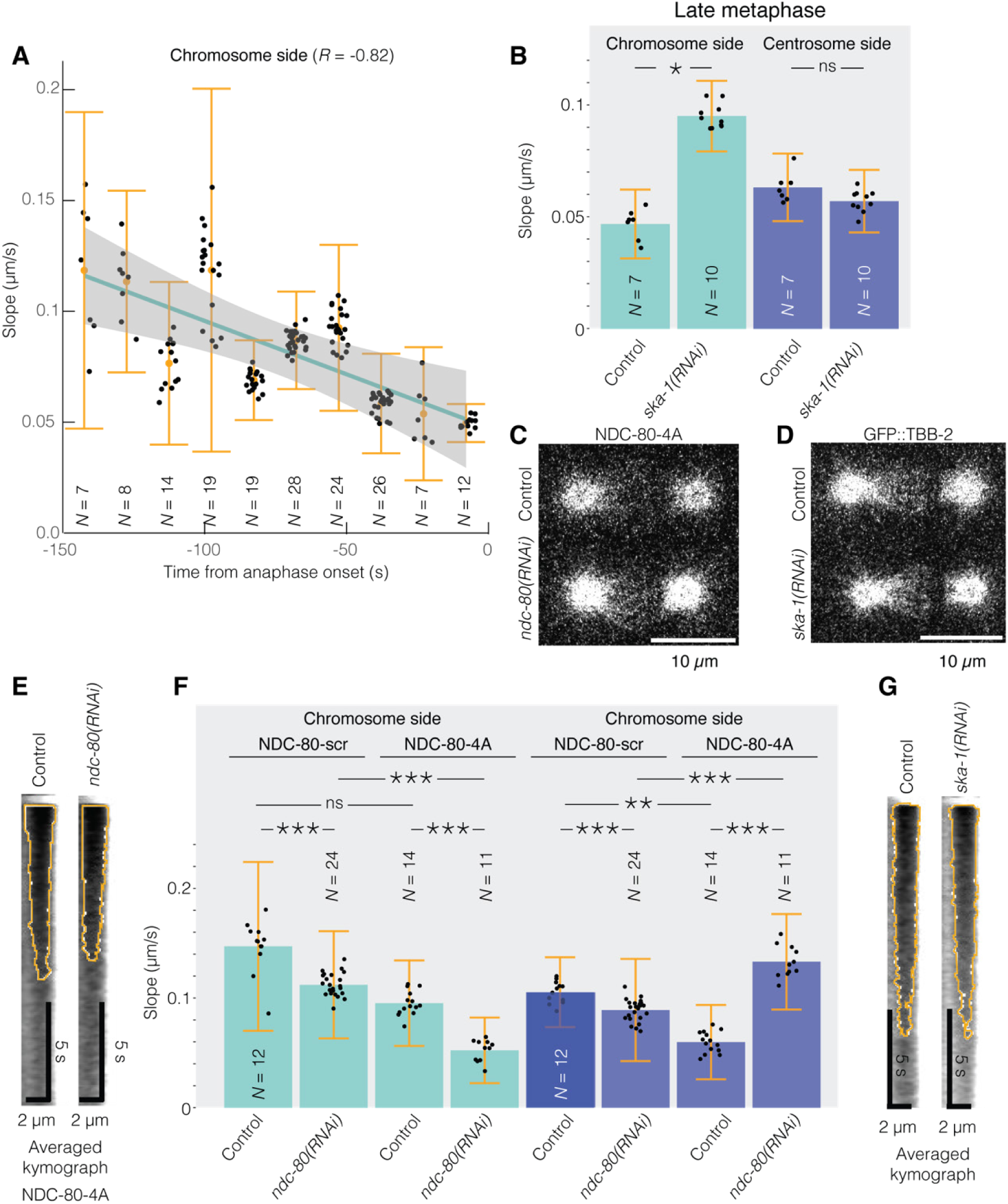
(**A**) Correlation between the time of bleaching and the front velocities by segmenting the dark regions of non-treated or control embryos of experiments reported in other figures. Microtubules were labelled using GFP::TBB-2^β-tubulin^. Black dots represent averages of N-1 embryos, leaving out, in turn, each embryo (Methods § Statistics on kymograph front slopes). Error bars are estimated using Jackknife resampling. Pearson coefficient read *R* = -0.82 (*p* = 0.0037). The grey-shaded region corresponds to the confidence interval at 0.95 on the correlation-line coefficients. (**B**) Front velocities by segmenting the dark region of *N* = 7 *ska-1(RNAi)* embryos bleached during late metaphase (−60 to -30 s from anaphase onset) and their *N* = 10 controls. The experiment was replicated (Fig S9C). (**C**, **D**) Exemplar micrographs of single (C) NDC-80-4A GFP::TBB-2 or (D) GFP::TBB-2^β-tubulin^ embryos used for FRAP experiment and submitted to (C) *ndc-80(RNAi),* (D) *ska-1(RNAi)* or corresponding control treatments. (**E, G**) Averaged kymographs over the posterior half-spindle for the data presented in (F and B, respectively), with centrosome on the right-hand side. The orange line delineates the bleached region as obtained by our analysis (Methods § Image processing). (**F**) Front velocities by segmenting the dark region of NDC-80-scr embryos, either (*N* = 9) treated by *ndc-80(RNAi)* or (*N* = 7) control and bleached; NDC-80-4A embryos either (*N* = 6) treated by *ndc-80(RNAi)* or (*N* = 7) control. Bleaching was performed during late metaphase (−60 to -30 s from anaphase onset). In panels (B and C), microtubules were labelled using GFP::TBB-2^β-tubulin^. Black dots represent averages of N-1 embryos, leaving out, in turn, each embryo (Methods § Statistics on kymograph front slopes). Bars correspond to means; error bars are estimated standard errors using Jackknife resampling.

To rule out the possibility that a cell-scale change in microtubule dynamics over time accounted for the above results, we genetically manipulated kinetochore-microtubule dynamics while maintaining their attachments. We depleted SKA-1 protein by RNAi, preventing the locking of end-on attachments. In contrast to mammalian cells, it did not delay anaphase onset (Cheerambathur et al., 2017; Schmidt et al., 2012). We monitored spindle length during the interval 5 to 15 seconds before anaphase onset (Suppl Methods). Using the data of the *ska-1(RNAi)* experiment and its replica, we measured a longer spindle in the perturbed condition, 15.4±0.2 µm (*N* = 8) compared to 14.3 ± 0.1 (*N* = 15, *p =*0.01), as previously reported. We measured the velocity of the fronts in the time range from 60 to 30 s before anaphase onset and observed a rate faster by 103% in depleted embryos (Fig 4B, replicated in Fig S9C) while no such decrease was visible in the same condition 120 to 60 s before anaphase onset (Suppl Fig S9C). This is consistent with the recruitment of SKA-1 in late metaphase (Cheerambathur et al., 2017). It once again supported our hypothesis that the chromosome-side front motion is caused by kMT poleward flux.

We then performed the converse experiment, inducing precocious and stronger recruitment of the SKA complex, thereby decreasing kinetochore-microtubule dynamics. We used the previously published mutated *ndc-80*, in which four amino acids at Aurora kinase phosphorylation sites were changed to alanines and termed it NDC-80-4A (Cheerambathur et al., 2017). The authors made it RNAi-resistant by changing some triplets to synonymous codons. We also considered the strain, with similar scrambling of gene coding but no alteration of the phosphorylation sites, as a control, termed NDC-80-scr. It enabled us to target the endogenous copy by RNAi. We crossed these strains with the one carrying the GFP::TBB-2^β-tubulin^ labelling and measured flux in the 120-30 seconds before anaphase onset. Upon *ndc-80(RNAi)*, NDC-80-4A displayed a shorter spindle, 13.1 ± 0.3 µm (*N* =3) compared to 14.6 ± 0.4 µm (*N* = 15, *p* = 0.001) in the 15-5 seconds interval before anaphase onset. Comparing NDC-80-4A conditions to NDC-80-scr, we observed a decreased chromosome-side front velocity by 53% and 35% upon *ndc-80(RNAi)* or control RNAi, respectively. This was expected, considering that SKA decreased the kMT dynamics at the kinetochore too early and too strongly (Fig 4C). In contrast, on the centrosome side, upon depleting the endogenous NDC-80 protein and rescuing with the mutated protein NDC-80-4A, we observed an increase in front velocity, suggesting that the precocious recruitment of SKA affected the front on each side differentially.

Altogether, the temporal evolution of the velocity of the chromosome-side front and its dependence upon the SKA complex confirmed that the kinetochore microtubules undergoing a poleward flux likely contributed to the chromosome front motion.

### The tetrameric kinesin KLP-18 is likely sliding the kinetochore microtubules along the spindle microtubules

Which mechanism could account for the kMT poleward flux? The most classic one is treadmilling, powered by the depolymerisation at the minus end. Electron micrographs showed that only a fifth of the kMTs reached their corresponding poles (Redemann et al., 2017). Besides, half of the microtubules had open minus ends at the centrosomes (O’Toole et al., 2003). The treadmilling mechanism usually involves MCAK as the main microtubule depolymeriser (Arpag et al., 2020; Brust-Mascher et al., 2004). The MCAK homolog, KLP-7, is also localised at kinetochores and spindle poles (Han et al., 2015; Sarov et al., 2012). We depleted this protein by RNAi, partially to preserve a functional spindle, although leading to a shorter spindle: 12.61 ± 0.13 µm (*N* = 9, t-test *p* = 0.001) compared to 14.36 ± 0.36 (*N* = 9) in control RNAi embryos. We measured the front velocity again (Fig S10). We did not observe a decreased front velocity but rather a 29% increase on the chromosome side. The trend was instead an increase in velocity that we attributed to a role of KLP-7 at the kinetochore, putatively facilitating the turnover of the kMTs (Jaqaman et al., 2010; Kline-Smith et al., 2004; Wordeman et al., 2007).

*In vitro*, a minimal treadmilling system comprises CLASP^CLS-2^, XMAP215^ZYG-9^, MCAK^KLP-7^ and EB1^EBP-2^ (Arpag et al., 2020). We thus performed a similar FRAP experiment and analysis under the null-mutant condition *ebp-2(*gk756) (Fig. S11). We found a mild, although significant, reduction of the recovery-front rate on the chromosome side, by 15%. It was consistent with the lack of a strong phenotype previously reported in the embryo for EBP-2 depletion (Kamath et al., 2003; Sonnichsen et al., 2005). However, several proteins involved in the kinetochore may be affected by EBP-2 depletion, such as SKA-1 (Lange et al., 2019) and the dynactin subunit DNC-2^p-50^ (Barbosa et al., 2017). In conclusion, we considered treadmilling an unlikely mechanism. We rather suggested that the kMTs moved locally poleward in a mechanism reminiscent of the poleward flux despite these microtubules being too short to reach the centrosome.

As an alternative, we considered the sliding of the kMTs along the sMTs, which were not moving. KLP-18, a kinesin-12 motor, appeared as a plausible candidate. Its mammal counterpart, KIF15, was shown to slide parallel microtubules *in vitro* (Drechsler and McAinsh, 2016). Furthermore, KLP-18 is also essential to correctly align the microtubules in the acentrosomal meiotic spindle of the nematode (Cavin-Meza et al., 2022; Wolff et al., 2022). Importantly, KLP-18 is localised in the mitotic spindle (Suppl Video S2). We thus set out to deplete this protein by RNAi. Because it is required for successful meiosis, conditions were hypomorphic. We measured a modest, although significant, decrease by 14% in the chromosome-side front velocity. We however observed a similar effect on the centrosome side (Fig 5A).

**Figure 5:**
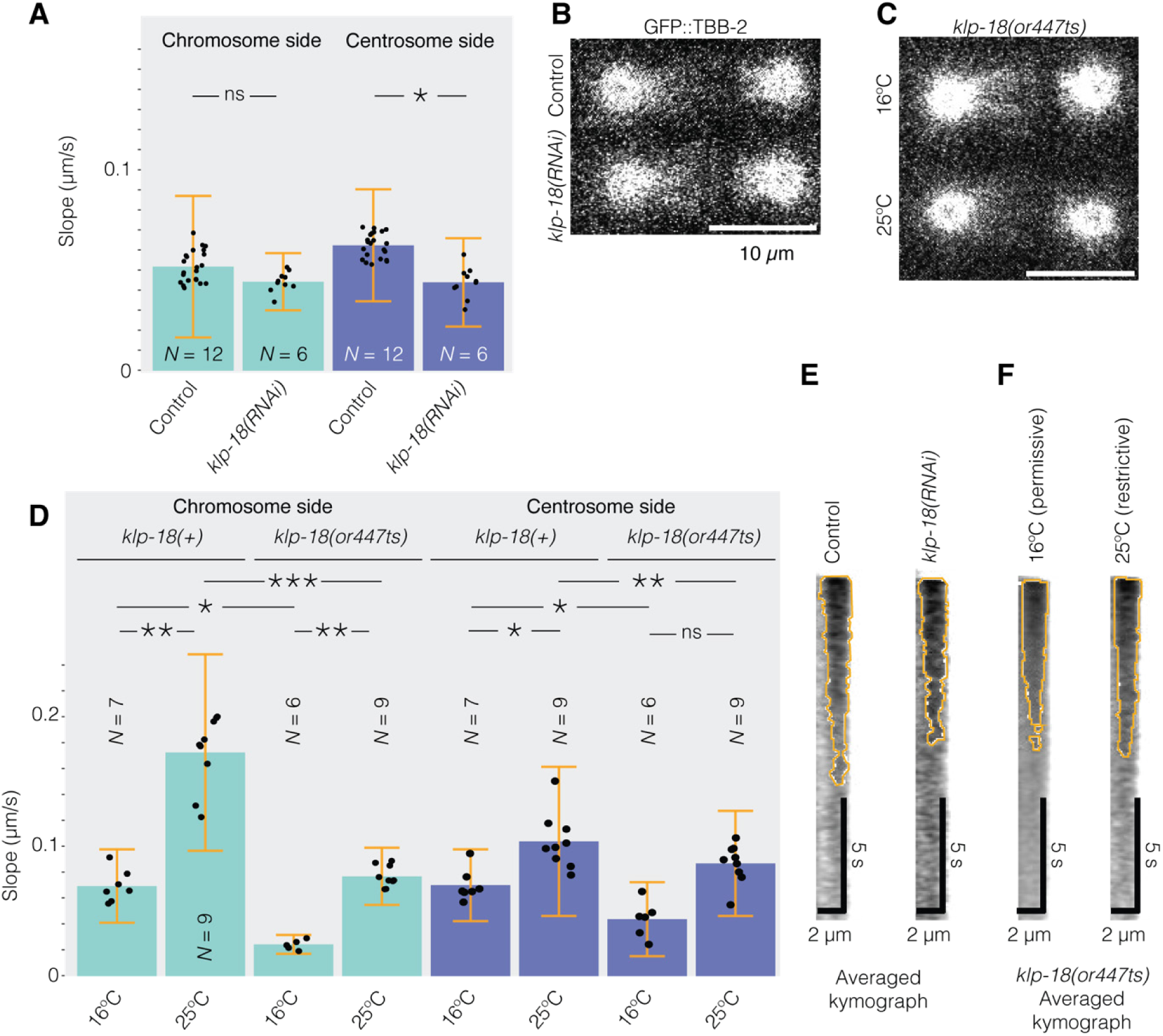
(**A**) Front velocities by segmenting the bleached region of *N =* 6 hypomorphic *klp-18(RNAi)* embryos compared to *N* = 12 control embryos. (**B**, **C**) Exemplar micrographs of single (B) GFP::TBB-2^β-tubulin^ or (C) *klp-18(or447ts)* GFP::TBB-2 HIS-11:mCherry embryos used for FRAP experiment and submitted to (B) *klp-18(RNAi)*, (C) restrictive temperature, or corresponding control treatments. (**D**) Front velocities by segmenting the bleached region of *klp-18(or447ts)* embryos at (*N* = 6) permissive temperature 16°C and (*N* = 9) restrictive temperature 25°C; corresponding control embryos klp-18(+) at (*N* = 7) permissive temperature 16°C and (*N* = 9) restrictive temperature 25°C. Microtubules were labelled using GFP::TBB-2^β-tubulin^. The experiment was replicated (Fig S12). (A, D) Black dots represent averages of *N*-1 embryos, leaving out, in turn, each embryo (Methods § Statistics on kymograph front slopes). Bars correspond to means; errors are estimated using Jackknife resampling. Light blue bars are values on the chromosome side and dark blue on the centrosome side. (**E, F**) Averaged kymographs over the posterior half-spindle for the data presented in (A and D, respectively), with centrosome on the right-hand side. The orange line delineates the bleached region as obtained by our analysis (Methods § Image processing).

To get a stronger phenotype, we considered the temperature-sensitive mutant *klp-18(or447ts)* compared to OD868 denoted *klp-18(+)* (Connolly et al., 2014). After culturing the worms at 15°C, they were either imaged at 15°C (permissive condition) or incubated at 25°C for 30 min prior to dissection and imaging (restrictive). With this treatment, we observed 4/4 visually normal meiosis at 16°C but 11/24 at 25°C. We selected embryos without defects in the number or appearance of pronuclei and measured the velocity of the front on the chromosome side. We observed a 114% decrease in embryos carrying the mutation compared to control embryos at the restrictive temperature (Fig 5B). It is noteworthy that the microtubule growth rate increases with temperature, accounting for the observed increase in front velocity between permissive and restrictive temperatures (Chaaban et al., 2018). In contrast, on the centrosome side at restrictive temperature, the velocity of the front decreased by 16%. We replicated this result (Suppl Fig S12). We concluded that KLP-18 specifically contributed to the front velocity on the chromosome side. We propose that this tetrameric kinesin slides the kinetochore microtubules along the spindle ones, remaining immobile.

## DISCUSSION

We investigated the dynamics of the microtubules within the metaphase spindle in the nematode one-cell embryo. While we confirmed the lack of significant global poleward flux by FRAP and photoconversion experiments, we observed a recovery of tubulin fluorescence within the bleached region that cannot be accounted for by the diffusion of tubulin dimers from the cytoplasm. Instead, we observed two opposing fronts closing the bleached region. The centrosome side front was accounted for by the combination of microtubule plus-end growth and diffraction in microscopy. Modelling such a behaviour confirmed that it causes a recovery of the bleached region, uniform along the spindle. Interestingly, such a mechanism is expected to act equally on both sides, and far from the chromosomes, where one sees only the spindle microtubules, we measured similar velocities on both sides. In contrast, and near the chromosomes, the front appeared to reflect a distinct mechanism superimposed on the previous one. We attributed it to the kinetochore microtubules sliding along the spindle microtubules, the latter being fixed. Reasons are fourfold: (i) We only saw the extra velocity due to kinetochore microtubules where they were in the majority over the other spindle microtubules, i.e. close to kinetochores. It was plainly consistent with the lack of flux detection in previous works (Labbe et al., 2004; Redemann et al., 2017). (ii) We measured a flow of microtubule minus-ends from the chromosomes towards the centrosomes, revealed by the optical flow of ASPM-1; (iii) A growth of the microtubules and a dynamic attachment at the kinetochores is needed to observe normal chromosome side recovery, as found by partially depleting ZYG-9^XMAP215^, NDC-80, or CLS-2^CLASP^. It is noteworthy that we also found accelerated flux upon *ndc-80(RNAi)*, *cls-2(RNAi)* and *ska-1(RNAi),* as expected for poleward flux in other organisms (Matos et al., 2009). (iv) Furthermore, the chromosome-side velocity depended on the status of the attachment of the microtubule at the kinetochore, either its normal changes over time, or altering it genetically. Indeed, we caused either premature or insufficient securing of the attachments by the SKA complex. Our suggested sliding mechanism contrasts with the previous proposal by Redemann and colleagues. Indeed, we found no indication that the kMTs are detaching from the centrosomes and moving towards the kinetochore. However, we cannot exclude the possibility that the kinetochore microtubules depolymerise from their minus-ends, consistent with 38% of the ends being opened in metaphase (Redemann et al., 2017).

Which mechanism could account for this sliding? A classic mechanism is treadmilling, requiring depolymerisation at the minus end. This mechanism usually involves MCAK as the main microtubule depolymeriser (Arpag et al., 2020; Brust-Mascher et al., 2004). In our case, because the depletion of the homolog of MCAK, KLP-7, did not decrease the recovery velocity but instead tended to increase it, we consider this mechanism unlikely. However, it is not impossible that other microtubule depolymerisers, like KLP-13, katanin, stathmin, and fidgetin, may contribute (Zhang et al., 2007). However, the depletion of KLP-13 has a reduced phenotype in the zygote (Kamath et al., 2003; Sonnichsen et al., 2005). Katanin was reported to be inactive during mitosis (Clandinin and Mains, 1993). Stathmin and fidgetin display a weak phenotype upon depletion in the one-cell embryo (Kamath et al., 2003; Lacroix et al., 2014; Lacroix et al., 2016; Sonnichsen et al., 2005). We also considered the transport of the kMTs by cross-linking to the sMTs unlikely as these latter are not undergoing poleward flux (Miyamoto et al., 2004; Steblyanko et al., 2020; Uteng et al., 2008).. We cannot exclude that bridging microtubules are involved provided they are short enough to remain in the central part of the spindle (Jagric et al., 2021), however, they do not appear numerous on electron micrographs (Redemann et al., 2017). We rather propose that the sliding of kMTs over the sMTs is achieved by the kinesin-12 KLP-18. This motor contributes to focusing and organising the meiotic-spindle microtubules (Cavin-Meza et al., 2022; Wolff et al., 2022). Its mammal counterpart, KIF15, was shown to slide parallel microtubules *in vitro* (Drechsler and McAinsh, 2016). Some redundant sliding mechanisms may also exist, such as the transport of the kMT minus ends to the poles by dynein using LIN-5^NuMA^ (Elting et al., 2014). In all cases, the sliding mechanisms must be coupled with the polymerisation of the microtubules at the kinetochore, like in other organisms, to maintain chromosome attachment (Barisic et al., 2021).

We wondered whether restricting the poleward flux to the kMTs is a peculiarity of nematodes. Metaphase is very brief in the nematode embryo, which prevents observing the spindle in a steady state. Furthermore, careful observation of the pole-to-pole distance during the metaphase shows a slow but constant length increase. This situation contrasts with the *Xenopus laevis* spindle prepared from extracts and at steady-state, where a gelification transition was observed (Dalton et al., 2022). A plausible and additional cause of this difference is the strong pulling forces exerted on the spindle poles by the cortical force generators (Grill et al., 2001; Grill et al., 2003; Labbe et al., 2004). Compared to other cells, these forces are very high and participate in spindle elongation. One can speculate that sliding kMTs along the sMTs may mechanically insulate the kinetochores from the poles. In early metaphase, when pulling is weak, the sliding mechanism could generate enough force to ensure a correct function of the spindle assembly checkpoint and subsequent correcting of chromosome mis-attachments. Oppositely, in late metaphase, the strong pulling could be tempered down by letting the kMTs slide along the sMTs, leading to slow spindle elongation. Indeed, while the spindle slowly elongates in the non-treated embryos at that stage, decreasing cortical forces by depleting GPR-1 and GPR-2 reduces this phenotype (Cheeseman et al., 2005; Lewellyn et al., 2010). Overall, such a mechanism may maintain some tension at the kinetochores and be compatible with the SAC and correction mechanism function. Another important difference is the centromere organisation. Nematode chromosomes are holocentric; therefore, the kMTs do not gather to form a K-fiber. However, a recent electron microscopy study suggests that enough space exists between microtubules in HeLa cells to accommodate molecular motors, making possible a mechanism similar to the one reported here (Kiewisz et al., 2022). Consistently, it was recently reported in human cells that the dynamics of microtubules along the spindle axis are not uniform (Conway et al., 2022; Risteski et al., 2022). If our proposed mechanism also applies to these cells, the minus-ends of the kMTs would also move towards the poles in a poleward flux. Consistently, half of the kMTs in HeLa cells do not reach the corresponding centrosomes (Kiewisz et al., 2022). This observation also suggests that a mechanism similar to the one proposed for the nematode may occur in mammalian cells.

## MATERIAL AND METHODS

### C. *elegans* strains

All *C. elegans* strains were maintained at 20°C and cultured using standard procedures (Brenner, 1974). *C. elegans* worms were grown on NGM plates seeded with OP50 *E. coli* strain. *C. elegans* worm strains and genotypes are listed in Table 1.

**Table 1:**
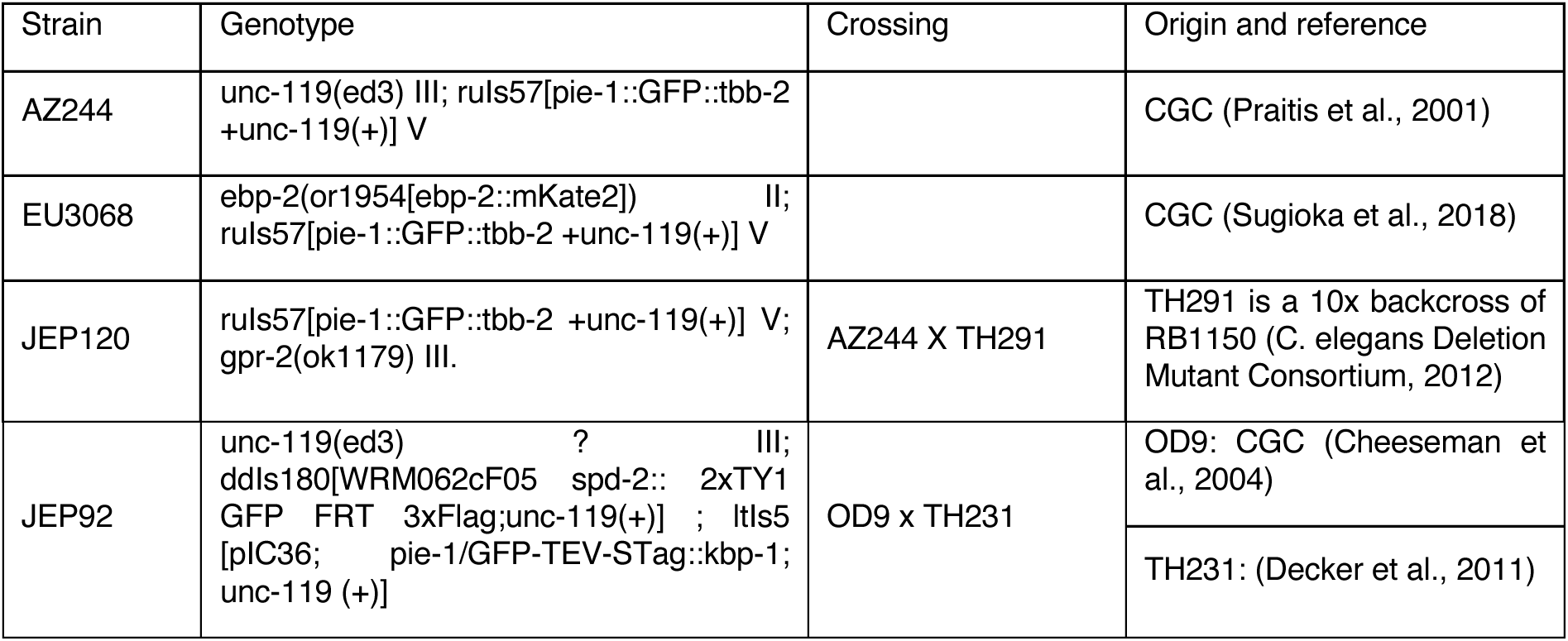
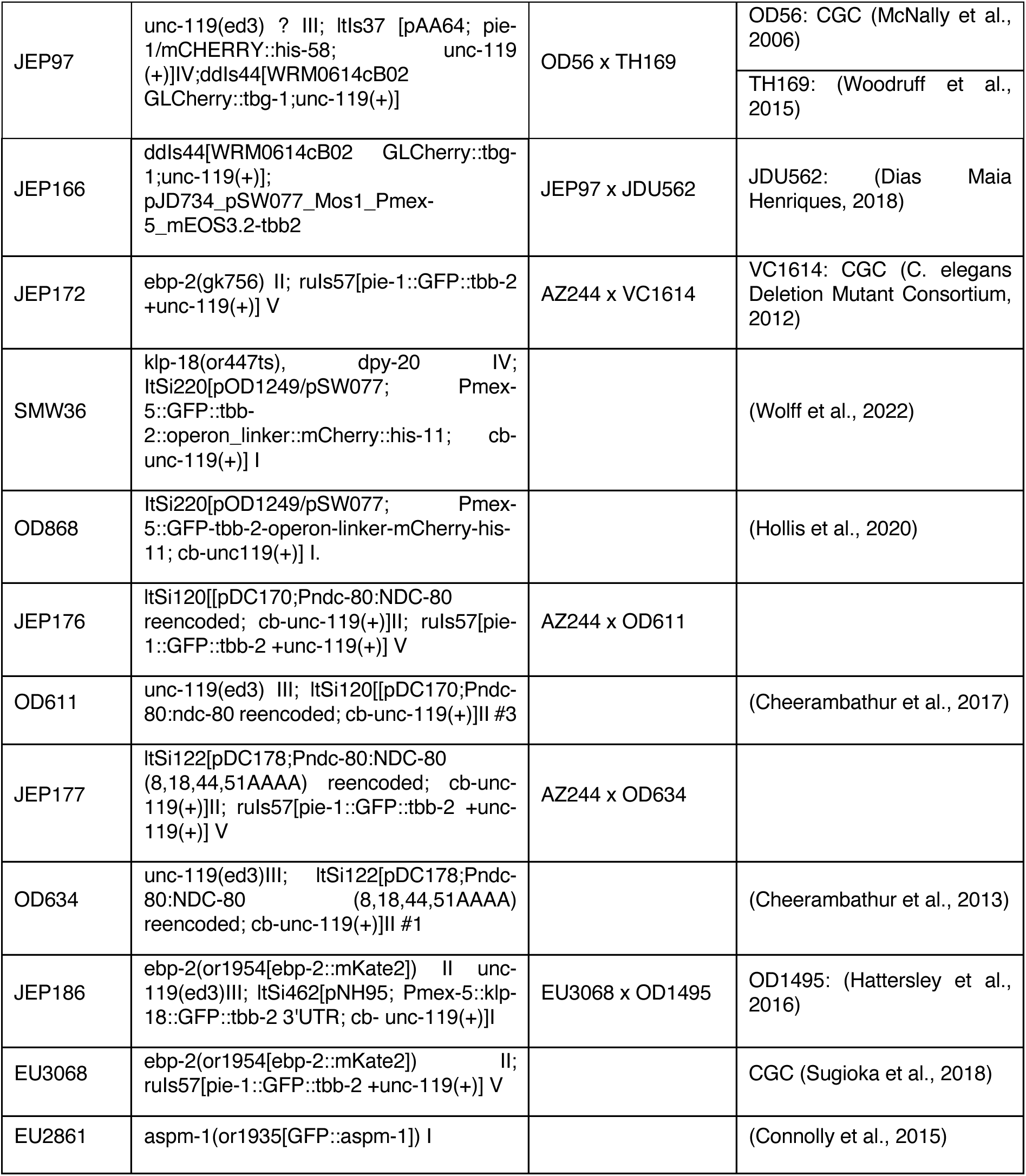
Worm strains used in this article and their genotypes.

### RNAi treatment

RNAi was performed by feeding as described previously (Kamath and Ahringer, 2003; Timmons and Fire, 1998). The control embryos for the RNAi experiments were fed with bacteria carrying the empty plasmid L4440. We targeted *ndc-80* by RNAi using the same sequence as in (Cheerambathur et al., 2017) after cloning it into a vector. All other clones came from the Ahringer-Source BioScience library (Kamath et al., 2003). RNAi target, clone ID or sequence used, and feeding conditions are listed in Table 2. We set the RNAi treatment duration so that we did not notice any phenotype suggesting impaired meiosis.

**Table 2:**
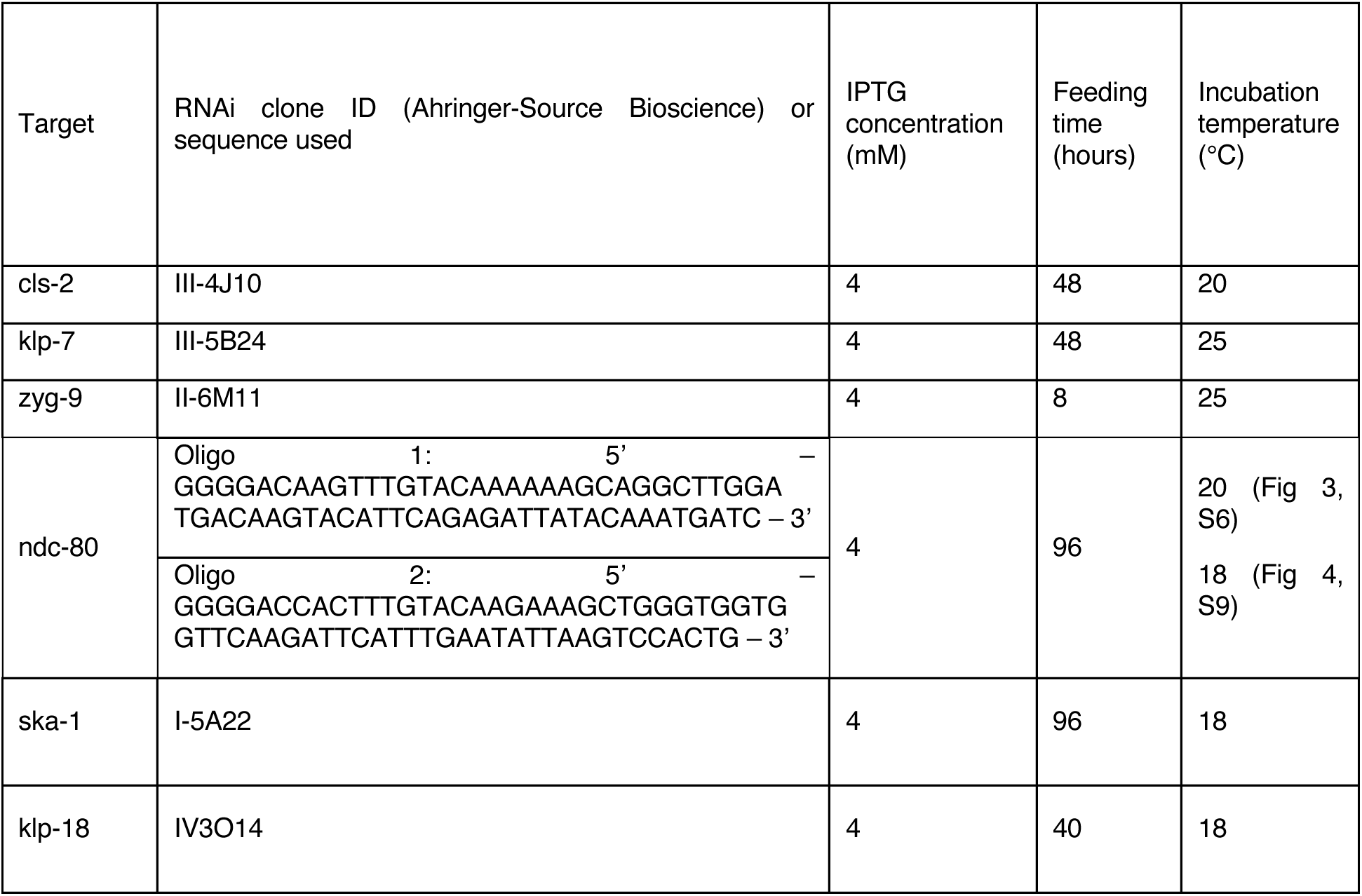
RNAi target, clone ID and treatment conditions used in this article.

### Microscopy: acquisition conditions

#### Preparation of C. elegans samples for imaging

*C. elegans* hermaphrodite adults were dissected in M9 medium (50 mM Na_2_HPO_4_, 17 mM KH_2_PO_4_, 86 mM NaCl, 1 mM MgSO_4_) (Brenner, 1974). The released embryos were deposited on an agarose pad (2% w/v agarose, 0.6% w/v NaCl, 4% w/v sucrose) between slide and coverslip or between two 24×60 cm coverslips. To confirm the absence of phototoxicity and photodamage, we checked for normal rates of subsequent divisions (Riddle, 1997; Tinevez et al., 2012).

#### Imaging condition for photobleaching and photoconversion experiments

Embryo imaging to perform FRAP was achieved on a scanning confocal microscope with deconvolution (LSM 880 airyscan, Zeiss) using an apochromatic 63x/NA1.4 oil objective. Acquisition parameters were controlled by Zen black software. We imaged at 18°C, unless otherwise stated, using the CherryTemp temperature control system (CherryBiotech, Rennes, France) in all cases. With all strains expressing GFP::TBB-2^β-tubulin^, we performed photobleaching using an argon laser at a wavelength of 488 nm and 70 µW power for 75 iterations (photobleaching time ∼ 1.9 s). We then imaged using the same laser at 2.30 µW. Five images were taken before photobleaching to obtain the embryo’s basal fluorescence. The laser power was measured at the objective output just before each microscopy session. We bleached a 2.6 x 19.5 µm area within the mitotic spindles (the embryo was oriented horizontally) (Fig1A). When accommodating two bleached regions per half-spindle (Fig 3AB), their width read 1.3 µm. When comparing recovery within the spindle and in the cytoplasm, we used a 2.6 x 9.5 µm area to be more selective and a frame rate of 1.5Hz. In all other cases, images were acquired at 12.5 Hz.

In photoconversion experiments, we used the strain JEP166 expressing mCherry::TBG-1^γ-tubulin^ and mEOS3.2::TBB-2^β-tubulin^. The mEOS3.2::TBB-2^β-tubulin^ labelling was obtained from the JDU562 strain, a kind gift from Julien Dumont’s team. The mCherry::TBG-1^γ-tubulin^ channel enables the registration of the images on the centrosomes’ position (see Methods § Image processing). Five images were taken before photoconversion to capture the embryo’s basal fluorescence. Photoconversion was performed using a 405 nm UV laser at 10 µW for 35 iterations (photoconversion time ∼ 0.850 s). We photoconverted a 13 x 1.3 µm area within the mitotic spindle after vertically orienting the anteroposterior axis of the embryo. Images were acquired with a 561 nm laser at 9 µW power and at 3 frames per second.

To study the localisation of KLP-18, we imaged *C. elegans* one-cell embryos at the spindle plane, from nuclear envelope breakdown (NEBD) through late anaphase. We used a Leica DMi8 spinning disk microscope with Adaptive Focus Control (AFC) and an HCX Plan Apo 100x/1.4 NA oil objective. Illumination was performed with a 488 nm laser, and we used a GFP/FITC 4 nm bandpass excitation filter and a Quad Dichroic emission filter. Images were acquired using an ultra-sensitive Roper Evolve EMCCD camera that was controlled by Inscoper, SAS acquisition software. During the experiments, the embryos were kept at about 20°C.

Finally, to investigate the localisation of ASPM-1::GFP, EBP-2::mKate2 or GFP::TBB-2 in control experiments, we imaged *C. elegans* one-cell embryos at the spindle plane during metaphase, on a scanning confocal microscope with deconvolution (LSM 880 airyscan, Zeiss) using an apochromatic 63x/NA1.4 oil objective. Temperature was maintained at 18°C using the CherryTemp temperature control system in all cases (CherryBiotech, Rennes, France). Acquisition parameters were controlled by Zen black software. Frame rate varied between experiment from 2.1 Hz to 6.3 Hz, and pixel size was either 130 nm or 170 nm.

In all experiments, the anaphase onset was taken as a time reference. Images were stored using OMERO software (Li et al., 2016).

### Image Processing: Automatic recognition of the bleached region edges and measuring the slopes of the fluorescence-recovery fronts

#### Assembling a homogenous set of embryos

Observing the kymograph (Fig 1D, S3) suggested that the recovery proceeded by two fronts moving inwards. We set out to measure the velocity of these fronts. We ensured the laser power remained the same at each session to safeguard against intensity variation. It also enabled us to have a set of embryos bleached in similar conditions. However, the spindle lies 10-15 µm deep in the sample, introducing some variability. Overall, we considered only the experiments where the FRAP efficacy reached at least 30%, assessed by comparing the fluorescence intensities before and after the FRAP event. We checked that we performed bleaching during metaphase, i.e. between - 120 and -30 s before the anaphase onset; it ensured a full fluorescence recovery prior to anaphase. For the *ska-1(RNAi)* experiments, this range was reduced to [-60 -30] s to correspond to the time period when SKA-1 is recruited in non-treated embryos (Cheerambathur et al., 2017). Finally, the distance between the metaphasic plate and the bleached region was measured to guarantee that a bright corridor separated the bleached region from the hollow due to the chromosomes, ensuring a robust measure of the chromosomal front speed (Fig 1B, S3).

#### Obtaining a kymograph for each embryo

We performed rigid registration of the raw images using a slightly modified version of the plugin (publicly available, v. 1.4) in Icy (de Chaumont et al., 2012). This registration focused on the centrosome position on the side of the bleached half-spindle, preventing spindle displacement or rotation from contributing to the measured front motion (Fig 1, S1, S2A). While we registered the raw images, we computed the rigid transform on a pre-processed image: we first deconvolved the microtubule images when focusing on anaphase because of the spindle rocking using DeconvolutionLab2 plugin (Sage et al., 2017); in all cases, we applied contrast-limited adaptive histogram equalisation (Zuiderveld, 1994). On the registered raw images, we applied a median filter under Fiji to limit the noise (Schindelin et al., 2012). We computed the kymograph by considering a region of interest of a length of 23.4 µm and a width of 3.12 µm (Fig S2B). To do so, we used Fiji and performed a median projection along a direction transverse to the spindle axis (Fig S2C).

#### Averaging kymographs over several embryos

To ensure that each embryo contributed equally, we performed a histogram equalisation among the individual embryos’ kymographs using the histogramMatcher class in sciJava (interfaced using Beanshell) (Rueden et al., 2021). Finally, we registered the kymographs aligning on the centrosome-side edge of the bleached region at the first time after bleach, using a supervised home-designed Matlab plugin (Fig S2C, arrowhead). We then averaged the kymographs pixel-wise (Fig S2D). We then cropped the embryo-averaged kymograph to consider only the first 40 seconds and the bleached half-spindle.

#### Kymograph segmentation

Next, we segmented the bleached area in the kymograph. As a preprocessing step, we convolved the kymograph with a 3×3 kernel that set the central element to 5 and the others to 1 (Gaussian blur). We then autonomously detected the non-bleached area by being conservative and considering only the pixels that were evidently non-bleached. To do so, we set the background class as the 60% brightest pixels. We then performed grey morphology (to remove micro/dot regions due to noise). Practically, we used two iterations of opening with a 1-pixel (130 nm) circular structural element and two closing iterations with the same element (Fig S2E, black line). Similarly, we set the bleached area as the 25% darkest pixels and performed the grey morphology (Fig S2E, orange line). We used neighbourhood information to decide about the pixels with intermediate grey levels. We trained a machine learning algorithm, namely random forests featuring 100 trees (Breiman, 2001), using the above defined classes for training across all kymographs in a set acquired under the same condition. We then applied the algorithm to classify the pixels within the kymograph (Fig S2E, red line) using Weka under Fiji (Beanshell script) (Arganda-Carreras et al., 2017). Finally, we performed grey morphology closing (2 pixels circle), watershed and the same closing again. It enabled us to remove artefactual dotty regions while preserving the largest one corresponding to the bleached area. We then selected the largest segmented object. Indeed, because the recovery was never complete (Fig 1D) and the contrast was low, a small artefactual region may appear in the tail of the main one. Finally, we dilated the shape using a 1-pixel-radius circle twice, making the resulting shape convex.

#### Kymograph boundary fitting

Having segmented the bleached area, we fitted a line along the boundaries (between 2 and 30 s) to get the mean front velocities. We also computed the mid-curve (average between the positions of the two edges at each time point) and fitted it with a line (Fig S2F).

#### Statistics on kymograph front slopes

We used a resampling method to estimate the error bars for the front velocities, since we could not compute them for individual embryos due to image noise. We used the Jackknife method, which requires computing the slope over a set containing all embryos but one (Efron and Tibshirani, 1993; Quenouille, 1956). We thus obtained a standard error for the averaged value estimated across all embryos and could perform *t*-tests against the null hypothesis that the slope is 0.

#### Statistics

To compare conditions, we used the Jackknife again (Arvesen, 1969; Schechtman and Wang, 2004) and performed a t-test on pseudo-values after (Tukey, 1958) or Pearson correlation when relevant. For the sake of simplicity, we depicted confidence levels using diamonds or stars (***, P ≤ 0.0005; **, P ≤ 0.005; *, P ≤ 0.05; ÷ P ≤ 0.1; n.s., P >0.1). This latter term means ‘non-significant’ and was often omitted for the sake of clarity.

## Acknowledgements

The strains TH291, TH231, and TH169 are kind gifts from Prof A. A. Hyman. Dr J. Dumont kindly offered JDU562. We thank Dr Gregoire Michaux for the feeding clone library and technical support. We are also thankful to Dr Arshad Desai for the gift of strains OD868, OD611, OD1495 and OD634. The strain SMW36 was a kind gift of Prof Sarah M. Wignall. EU3068 was a kind gift from Prof Bruce Bowerman. We also thank Drs. Gilliane Maton, Fabrice Mahé, Anne Corlu, Christophe Heligon, Rebecca Smith, Sebastien Huet, Flora Demouchy, Gregoire Michaux, Anne Pacquelet, Stéphanie Dutertre, Xavier Pinson, Marc Tramier, Ostiane d’Augustin for discussions about the project. Some strains were provided by the Caenorhabditis Genetics Center (CGC), funded by the National Institutes of Health Office of Research Infrastructure Programs (P40 OD010440; University of Minnesota). We also acknowledge La Ligue contre le cancer (comites d’Ille-et-Vilaine et du Maine-et-Loire). Microscopy imaging was performed at the Microscopy Rennes Imaging Center (MRIC), UMS 3480 CNRS/US 18 INSERM/University of Rennes 1. MDS acknowledge the fellowship by Fondation ARC (ARCDOC42024120009129). MRIC is a member of the national infrastructure France-BioImaging, supported by the French National Research Agency (ANR-10-INBS-04). In particular, we acknowledge the funding of the Zeiss Airyscan confocal microscope by EU funding FEDER under reference CARE Phase 2.

## Author contribution

Conceptualisation: JP, LLM; Data curation: NS, MDS, CT, LC, JP; Formal analysis: NS, MDS, CT, LC, JP; Funding acquisition: JP, HB, LLM; Investigation: NS, MDS, CT, PF, LC, SP, JP; Methodology: NS, MDS, HB, LC, JP; Project administration: JP; Software: NS, JP, HB; Supervision: LLM, JP, LC; Validation: HB, LC, JP; Visualisation: NS, JP; Writing NS, JP; Writing – review and editing: LC, NS, MDS, JP, LLM, HB.

