## Supplemental Text, Methods, Figures and Legends for "Kinesin-12 KLP-18 contributes to the kinetochore-microtubule poleward flux during the metaphase of *C. elegans* one-cell embryo"

We detail here complementary investigations and the approaches developed specifically for this work, while the details for applying existing methods are reported in the Supplementary Methods section below.

##### 1. Cytoplasmic tubulin-dimer diffusion does not account for recovery after photobleaching.

During metaphase, we bleached the fluorescently labelled tubulin in a band-shaped region, 2.6 μm-wide, perpendicular to the spindle axis and covering the spindle but extending up to the cytoplasm (Fig 1A); for analysis, this region was then broken up into two, one in the spindle and one in the cytoplasm, for analysis. The proximal boundary of the bleached area was between 0.5 and 2 μm away from the chromosomes (defined as the area devoid of microtubules in the micrograph). Since the spindle is moving and slowly elongating during metaphase, we performed an intensity-based registration of the images over time using the Icy software (de Chaumont et al., 2012). We used the centrosome on the bleached side as a spatial reference (see Methods § Image processing, Fig 1A). We monitored the fluorescence recovery over time and, assuming a single microtubule population, fitted the time-recovery curve with a single exponential over a 60-second range during metaphase (Giakoumakis et al., 2017; Girao and Maiato, 2020) (Methods § Microscopy). We used a global fitting approach to safeguard against variability between embryos and obtained the confidence interval by thresholding the empirical likelihood (Bouvrais et al., 2021). In doing so, we imposed the same model parameters on each embryo of a given dataset and maximised the product of the embryo-wise likelihoods (Beechem, 1992). We found a recovery half-time inside the spindle about three times longer than in the cytoplasm, although the recovery was never complete (Fig 1B). The spindle half-life was consistent with previous measurements (Redemann et al., 2017) and suggested that free fluorescent tubulin was available within seconds within the spindle.

##### 2. Bleached-area edge closure depends on microtubule dynamics.

To challenge the link between the observed fronts (Fig 1D) and microtubule dynamics, we sought to decrease microtubule growth rates globally. ZYG-9$^{XMAP215}$ was a suitable candidate, as it affects all microtubules, including astral, spindle, and kinetochore-associated ones (Encalada et al., 2005; Fernandez et al., 2009; Lacroix et al., 2018; Srayko et al., 2005). However, we restricted ourselves to a partial depletion to preserve functional spindle assembly (Matthews et al., 1998). We observed a delayed spindle rotation, confirming the penetrance of the treatment (Bellanger et al., 2007). We then performed the FRAP experiments and measured the absolute value of the slopes of the fronts (Fig S1B, C, replica in S1D). We observed a significant decrease in slope on both the chromosome and centrosome sides. It suggested that microtubule dynamics contributed to the mechanisms accounting for these fronts.

#### 3. Modelling recovery on the centrosome side

We reckoned that the front displacement on the centrosome side might be accounted for by the dynamics of the sMTs (growth and shrinkage) combined with diffraction due to imaging. Indeed, the fast recovery after photobleaching within the cytoplasm suggested that non-bleached tubulin diffuses fast and supports the growth of microtubules with bright dimers in the bleached area. We set out to model the kymograph by combining known microtubule dynamics and simulating the diffraction due to the microscope.

**Spindle- and kinetochore-microtubule dynamics**

We considered two microtubule populations within the spindle: one emanating from the centrosome and one attached to the kinetochore. In doing so, our analysis ignored the long-lived microtubules. These microtubules could be the few kMTs directly connecting the centrosome to the kinetochores, but their proportion is low (Redemann et al., 2017). Long-lived microtubules could also correspond to overlapping sMT. However, these latter were mostly expected around the metaphasic plate as it was suggested that microtubules could hardly perforate the chromosome domain (Redemann et al., 2017; Schneider et al., 2022). The region of interest used in extracting the experimental kymograph did not include many of these, particularly when close to the kinetochores. We thus ignored long-lived microtubules in modelling and focused on the first 30 seconds after photobleaching.

We modelled the spindle microtubules (sMTs) as free-growing microtubules until reaching the chromosomes (Zelinski et al., 2012) (Suppl File S1). Microtubule dynamics parameters were obtained from the literature or our measurements (Suppl Table S1). It resulted in an exponential-like decay of microtubule density from the centrosome to the kinetochores, consistent with electronic microscopy imaging (Fig S5A, red line) (Redemann et al., 2017). In particular, since the growth rate is critical to our model, we measured it to be $0.98 \pm 0.02$ μm/s (average and standard deviation of the Gaussian) under our conditions (Suppl Text §4, Fig S13). At the kinetochores or chromosomes, modelled as impassable obstacles (Schneider et al., 2022), we set the catastrophe rate to 1 per second. Indeed, varying by a factor of 10 around this value suggested that the density profile of the sMTs was relatively insensitive to this parameter. Furthermore, such a rate corresponded to a 1s residency time before detaching from the kinetochore and was similar to observations on astral microtubules at the cell cortex (Bouvrais et al., 2021). This catastrophe rate was about 5 times higher than at free microtubule ends. We adjusted the spindle length as the distance between the two fluorescent centrosomes in the images. We computed it using the intensity profile of non-treated embryos, averaged over 0.5 seconds, just after photobleaching. Technically, we smoothed the intensity profile using a cubic spline and set the positions of the peaks (assumed to correspond to the centrosomes) to the zero-crossings of the derivative.

We next modelled the kinetochore microtubules. We assumed that microtubules were permanently growing at kinetochores since they are under tension, as reported in other organisms (Cheeseman et al., 2004; Suzuki et al., 2015). The corresponding speed was set close to our observation, 0.1 μm/s. Doing so, we obtained kMT minus ends uniformly distributed along the spindle, as expected (Fig S5E, black line) (Redemann et al., 2017). We set the ratio of kMT to sMT to 1.5 at 1 μm from the kinetochore after the electron micrograph (Fig S5E).

**Recovery after Photobleaching**

We then modelled the fluorescence recovery after photobleaching. We assumed no exchange of tubulin along the lattice, i.e. no repair of microtubule defects (Schaedel et al., 2019). Recovery of the bleached region, where sMTs were in the majority, was attributed to microtubule dynamic instability. Technically, we modelled bleaching as the complete disappearance of fluorescence in the corresponding region. About recovery, based on the fast diffusion of tubulin dimers in the cytoplasm, we assumed that fluorescent tubulin dimers are readily available and modelled the growth of bleached microtubules with fluorescent subunits. It was equivalent to scaling the fluorescence curve in the bleached region with a coefficient (Fig S5A). This coefficient increased from 0 at the bleaching time to 1 upon reaching total recovery. We modelled the evolution of this coefficient using a single exponential function with a characteristic time of 30.4 s, as measured in this study. In doing so, we assumed that only free diffusion was involved. In the region studied, sMTs were the majority in most areas. Recovery on the kinetochore side was again attributed to the dynamic instability mechanism, but with kMTs poleward flux superimposed (Fig S5A). The flux rate was adjusted so that the velocity at the edge of the simulated bleached region mimicked the experimental value in non-treated embryos. Finally, we formed a pseudo-kymograph (Fig S5D).

##### Diffraction due to microscope imaging

We modelled diffraction using a Gaussian PSF, whose standard deviation was obtained by fitting the experimentally measured PSF and was read as 149 nm. To do so, we imaged fluorescent beads of diameters 175 nm (PS-Speck Microscope Point Source) on the same microscope. We fitted the experimental image with a Gaussian to obtain the PSF (Kirshner et al., 2013). We then convolved such a Gaussian, in 1D, with the simulated microtubule density at each time point (Fig S5BC) to obtain a convolved kymograph that mimics the experimental kymograph (Fig. 2A, compared with Fig. 1D and S5D). Notably, it differed from the one in Fig S5D, which did not account for microscopy diffraction.

##### Segmentation of the pseudo-kymograph and velocity extraction

We analysed this simulated kymograph (pseudo-kymograph) in a manner similar to the experimental one to extract the slopes of the fronts (Methods § Kymograph boundaries fitting). One parameter was changed: we used a 13% threshold on the intensity histogram to segment the bleached area. Indeed, the absolute intensity in the simulation differed from that in the experiment. This first segmented region, the core of the bleached region, was used to train the machine learning. We then performed the remaining steps of segmentation and kymograph edge fitting, as in the experimental kymographs. We obtained a front velocity of 0.016 µm/s on the centrosome side, in the same order of magnitude as the experiments. The bare growth and shrinking of microtubules emanating from the centrosome (Fig S5A), combined with diffraction due to microscope imaging, were sufficient to produce an apparent front movement. It could account for the recovery observed on the centrosome side, despite the sMTs not undergoing flux (Fig S5B). When no kMT is added, the front motions, without flux, were predicted to be almost identical on both sides. Adding fluxing kinetochore microtubules broke this symmetry in agreement with experiments (Fig S5C).

| Quantity | Value | Reference |
|---|---|---|
| MT growth rate | 0.65 µm/s | (Srayko et al., 2005) |
| MT shrinking rate | 0.84 µm/s | (Kozlowski et al., 2007) |
| Catastrophe rate | 0.28 /s | (Lacroix et al., 2018) |
| Rescue rate | 0.44 /s | (Lacroix et al., 2018) |
| Catastrophe against Chromosome | 1 /s | (Bouvrais et al., 2021) |
| Spindle length | 14.5 µm | This study |
| kMT poleward displacement | 0.1 µm/s | This study |
| sMT recovery characteristic time | 30.4 s | This study |

| | | |
|---|---|---|
| Bleached region boundary from the centrosome | 3.18 $\mu$m | This study |
| Bleached region width | 2.6 $\mu$m | This study |
| PSF standard deviation | 149 nm | This study |

**Supplemental Table S1**: Values used in the simulation and associated references.

##### Validating the model

We set out to experimentally challenge the proposed model. We predicted that the slope on the centrosome side scaled inversely with spindle length (Fig 2C). We tested the correlation between the spindle length and the front velocity on the centrosome side in control and non-treated conditions used in this article (Fig 2B). We observed an anti-correlation (Pearson $r$ = -0.82, $p$ = 0.09) as predicted by the model (Fig 2C). Interestingly, such an anti-correlation was not observed experimentally for the front velocity on the chromosome side (Fig S4). It is consistent with the model prediction (Fig 2C). We concluded that the front motion closing the bleached area on the centrosome side was likely not caused by any microtubule flux. Instead, it was due to the combination of spindle microtubule dynamics and microscopy imaging.

#### 4. Measuring the growing dynamics of spindle microtubules

We measured the displacement of the microtubule plus-ends within the spindle using EBP-2$^{EB1}$ labelling (Srayko et al., 2005). We imaged the doubly labelled strain EBP-2$^{EB1}$::mKate2; GFP::TBB-2$^{\beta\text{-tubulin}}$ at two frames per second to maintain a high SNR. We registered these images over time to suppress the effect of spindle displacement (Methods § Image processing), then denoised them with the Kalman filter (Kalman, 1960) (Fig S13A) and formed the kymograph as previously described (Fig S13B). We then measured microtubule plus-end velocity using the directionality ImageJ plugin in Fiji (Suppl Methods § Measuring the growth rate of microtubule plus ends), disregarding whether this velocity was due to microtubule growth or displacement (Liu, 1991; Schindelin et al., 2012; Schneider et al., 2012). We noticed that the distribution of comet orientation angles for a single embryo corresponded to a single Gaussian (S13D). We thus performed a global fit of individual embryos transforming angles into velocities (equation in Methods § Measuring the growth rate of microtubule plus ends) and found a growth rate of 0.98 ± 0.02 µm/s (average and standard deviation of the Gaussian), consistent with previous measurements (Lacroix et al., 2018; Srayko et al., 2005). The histogram of EBP-2$^{EB1}$ comet speed is reproduced in Fig S13C. It suggested that a single dynamical behaviour was present far from the chromosomes (where we measured), likely corresponding to the growing sMTs.

#### 5. Alternative measure of microtubule displacement towards the poles.

To ascertain our measurement of the poleward flux of the kinetochore microtubule, we sought an alternative method. We imaged the mitotic spindle with a confocal microscope, focusing on the microtubule minus-ends. To do so, we used ASPM-1 labelled with GFP, which marks the poles of the female meiotic acentrosomal spindle (Connolly et al., 2015). Indeed, the human homolog, ASPM1, was localised to the microtubule minus ends *in vitro* (Jiang et al., 2017; Mullen and Wignall, 2017), while C. *elegans* ASPM-1 was consistently seen concentrated at the spindle poles in meiosis (Connolly et al., 2015; Ellefson and McNally, 2011; Harvey et al., 2023; Taylor et al., 2023). We imaged this strain using a confocal microscope with deconvolution (Methods § Microscopy, Movie S3, Fig. S7A). Because the ASPM-1 spots were too faint to be tracked, we analysed optical flow in the mitotic spindle halves using the approach published by Drechsler and colleagues (Drechsler et al., 2020). We registered the images on the posterior side of the spindle, as in the FRAP analysis, to align the spindle along the anteroposterior axis (Methods § Image Processing). We then delineated two 3.8x5.8 μm regions of interest covering the anterior and posterior halves of the spindle and analysed the optical flow within them. We considered only the last 30 seconds of metaphase.

The algorithm performed in two steps: first, denoising the image through a variational scheme regularised by limiting the spatial and temporal first derivatives, weighted by hyperparameters $\alpha_1$ and $\beta_1$, respectively. Then, from the denoised image, the displacement vector field is estimated through a second variational scheme regularised in space and time by hyperparameters $\alpha_2$ and $\beta_2$, respectively. Setting proper values was crucial for detecting optical flow. Therefore, we tested hyperparameters in the ranges: $3.5 \times 10^{-4} < \alpha_1 < 3.5$ ; $2.5 \times 10^{-2} < \beta_1 < 25$ ; $3.5 \times 10^{-9} < \alpha_2 < 3.4 \times 10^{-3}$ ; $\times 10^{-8} < \beta_2 < 4 \times 10^{-3}$. To seek optimal values, we observed the flow (a vector field of the same size as the image, at each time point) and broke it into a 5x5 tiling. We computed the Shannon entropy in the centre tile, which corresponds to the zygote cytoplasm including the spindle, and in the 10 peripheral tiles, which correspond to the media outside the embryo. We then computed the median of the tile-wise values obtained in the periphery at each time point. They will be further referred to as the centre and edge median instantaneous entropies. Finally, we took the median over time for each embryo, and then over embryos of the same condition. Values between the sampled ones were interpolated using linear radial basis functions with a second-order polynomial added. We formed a cost function as a weighted sum of the squared median-centre entropy and the squared median-edge entropy. We minimise it through sequential least-squares programming to get the optimal set of hyperparameters: we obtained $\alpha_1 \simeq 0.32$, $\beta_1 \simeq 0.04$, $\alpha_2 \simeq 5.8 \times 10^{-7}$, $\beta_2 \simeq 0.0035$. We noticed that $\alpha_2$ was critical. We manually screened this latter to obtain the optimal $\alpha_2 \simeq 2.9 \times 10^{-6}$.. With these hyperparameter values, we obtained an optical flow showing clear biases towards centripetal displacement around both anterior and posterior centrosomes (Fig S7EF).

To test the significance of this bias in optical flow, we observed that the flow direction at the same position in the flow field does not change rapidly over time. We therefore discretised the directions into six equal angular sectors. We focused on two classes: one corresponding to the (-30, 30) degree sector, termed the right direction, and one corresponding to the (150, -150) degree sector, termed the left direction. For each position in the flow field, we considered the temporal statistical mode, i.e., the most common direction. We then formed histograms of direction for the resulting time-mode flow field at various positions within the region of interest. On the anterior side, 84.9±11.4 % (mean ± SD, $p = 6.6\times10^{-4}$ t-test against 0.5, $N = 6$) of the positions display a centripetal flow oriented in left direction. On posterior, the proportion read 97.5 ± 2.1 % ($p = 3.5\times10^{-8}$) oriented in right direction. We concluded that ASPM-1 and thus the minus ends of the microtubules moved centripetally.

As a positive control, we performed a similar analysis, including the hyperparameter optimisation on the strain EU3068 carrying the labelling EBP-2::mKate2. We obtained $\alpha_1 \simeq 0.05$, $\beta_1 \simeq 0.15$, $\alpha_2 \simeq 2.4 \times 10^{-5}$, $\beta_2 \simeq 3.9$. This protein reveals the microtubules plus ends, and as expected, we measured a centrifugal optical flow (Srayko et al., 2005) (Fig S7GH). We measured a clear bias of plus-ends proportion moving centrifugally, 68.8 ± 29.2 % ($p = 0.031$, $N = 14$) in right direction on anterior side and 81.5 ± 16.8 % ($p = 9.0\times10^{-6}$) in left direction on the posterior.

Finally, as a negative control, we used the same strain and focused on GFP::TBB-2 labelling. Optimised hyperparameters read $\alpha_1 \simeq 0.16$, $\beta_1 \simeq 1.9$, $\alpha_2 \simeq 4.9 \times 10^{-7}$, $\beta_2 \simeq 0.067$. We observed no clear antero-posterior directionality, indicating that the algorithm is not hallucinating (Suppl Fig S7IJ). We measured 58.9 ± 15.0 % ($p = 0.053$, $N = 13$) of the pixels with centripetal flow (leftwards) and 41.1 ± 15.0 % centrifugal (rightwards) on the anterior spindle half. We also measured and 34.1 ± 10.1 % ($p = 0.0001$, $N = 13$) centripetal (rightwards) and 66.0 ± 10.0 %

centrifugal (leftwards) on the posterior one. It suggests no preferred directionality of tubulin optical flow.

We repeated this analysis with 4 and 10 sectors and got equivalent results. We concluded that at least a fraction of the microtubules had their minus ends within the spindle moving towards the poles, supporting the proposed mechanism based on FRAP experiments.

### Supplemental Methods

#### *Imaging condition for measuring the velocity of microtubule plus-ends*

Embryo imaging was performed on a scanning confocal microscope with deconvolution (LSM 880 airyscan, Zeiss) using an apochromatic 63x/NA1.4 oil objective. Acquisition parameters were controlled by Zen black software. Imaging temperature was controlled using the CherryTemp temperature control system (CherryBiotech, Rennes, France). We also used focus maintaining for this particular experiment to compensate for drift. We measured the comets' velocity of microtubule plus-end to assess the microtubule growth rate. We used the strain EU3068 expressing EBP-$2^{EB1}$::mKate2 GFP::TBB-$2^{\beta\text{-tubulin}}$ (Sugioka et al., 2018). Image acquisition was performed with a HeNe laser at wavelength 594 nm and 3.35 μW power. The laser power was measured at the objective output at the beginning of each microscopy session. Images were acquired at 2 Hz on a single plane.

#### *Measuring the growth rate of microtubule plus ends*

We measured the microtubule's growth rate within the spindle by looking at its plus-end displacement. First, we registered the images using the GFP::TBB-$2^{\beta\text{-tubulin}}$ channel to keep the posterior centrosome immobile in the image stack as described above (Methods § Averaging kymograph over several embryos) (Fig S13A). Indeed, our study of the flux focused on the spindle half. Using the Kalman filter (Fiji plugin *Kalman Stack Filter*, with *noise estimate* = 0.2 and *bias* = 0.8), we denoised the EBP-$2^{EB1}$::mKate2 channel (Kalman, 1960). The EBP-$2^{EB1}$ labelled plus-end displacement appeared as an oblique line on the kymograph (Fig S13B). This latter was obtained by considering a region of interest of a length of 4.68 μm and a width of 3.12 μm centred on the posterior half-spindle. We formed the kymograph as previously described. We measured the direction and slope of these lines using the slightly modified version of the directionality plugin on ImageJ ((Tinevez et al., 2017), https://imagej.net/plugins/directionality). It produced a histogram counting the lines per direction (Fig S13D). We modelled the orientation distribution with a Gaussian. $a_d.e^{-(x-\mu_d)^2/2\sigma_d^2}$ with $\mu_d$ the average orientation, $\sigma_d$ the standard deviation and $a_d$ the normalisation factor. We computed the velocity corresponding to each embryo's direction distribution. However, the focus-maintaining system caused variable frame rates. We thus performed a global fit over all embryos, sharing the parameters to safeguard embryo-to-embryo variability again (Beechem, 1992). Practically, we used the model $\hat{a}.\exp\left(-(\text{atan}(v.\frac{\tau}{\rho})-\text{atan}(\hat{\mu}.\frac{\tau}{\rho}))^2 \Big/ 2\text{atan}(\hat{\sigma}.\frac{\tau}{\rho})^2\right)$ with $\hat{a}$ the fit-estimated normalisation factor, $v$ the velocity, $\tau$ the cycle time at imaging (different for each embryo), $\rho$ the pixel size (resolution), $\hat{\mu}$ the estimated velocity and $\hat{\sigma}$ the standard deviation (Fig S13C).

#### *Measuring the spindle length*

The spindle length was measured manually on images taken just before bleaching, if we do not expect a phenotype at a specific point in metaphase, using the Fiji distribution of ImageJ. In the case of *ska-1(RNAi)* or NDC-80-4A, the SKA complex intervenes only in late metaphase (Cheerambathur et al., 2017). We thus tracked the spindle poles using our previously published software (Pecreaux et al., 2006) and considered for each embryo the median length between 15 and 5 seconds before anaphase onset. We then computed the mean and its standard error over the embryos. Statistical tests used a Student's t-test with the Welch-Satterthwaite correction for unequal variances.

### Supplemental Figures

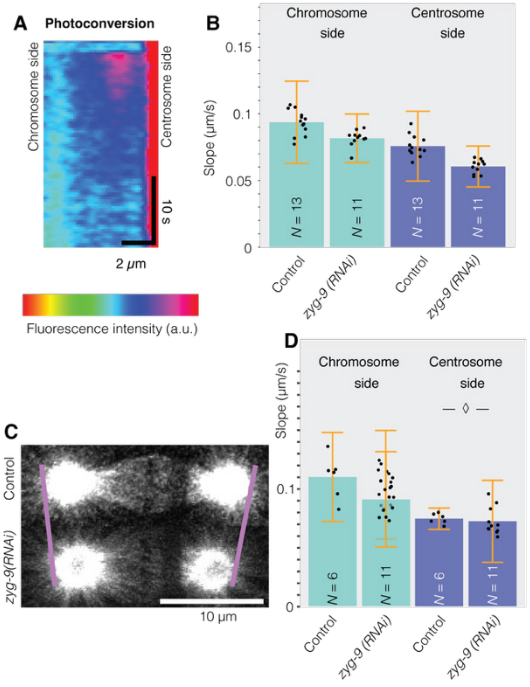


**Figure S1: Tubulin fluorescence recovery after photobleaching and photoconversion in the metaphasic spindle of *C. elegans*.** (**A**) kymographs after photoconversion within the mitotic spindle averaged over *N* = 7 dual labelling mEOS3.2::TBB-$2^{\beta\text{-tubulin}}$ ; mCherry::TBG-$1^{\gamma\text{-tubulin}}$ embryos. The first channel enabled visualising tubulin recovery, while the second enabled registration (Methods). The colour scale ranges from blue for dark pixels to red for bright areas. The centrosome was located on the right-hand side. (**B**) Front velocities by segmenting the bleached region of *N* = 11 *zyg-9(RNAi)* treated embryos and *N* = 13 control ones. Black dots represent the averages of *N*-1 embryos, leaving out, in turn, each embryo (Methods § Statistics on kymograph front slopes). Bars correspond to means; error bars are estimated standard errors using Jackknife resampling. Light blue bars are values on the chromosome side and dark blue on the centrosome side. Slopes are reduced by 13 ± 35% and 20 ± 34%, on the chromosome and centrosome side, respectively. (**C**) Exemplar micrographs of a single GFP::TBB-$2^{\beta\text{-tubulin}}$ embryo used for the FRAP experiment and submitted to (bottom) zyg-9*(RNAi)* or (top) corresponding control treatments. (**D**) Replica of the experiment (B) using *N* = 11 *zyg-9(RNAi)* treated embryos and *N* = 7 control ones. Light blue bars are velocity values measured on the chromosome side, and dark blue on the centrosome side. In B-D, we used strains with GFP::TBB-2β-tubulin-labelled microtubules. Slopes are reduced by 17 ± 46% and 3 ± 48%, on the chromosome and centrosome side, respectively.

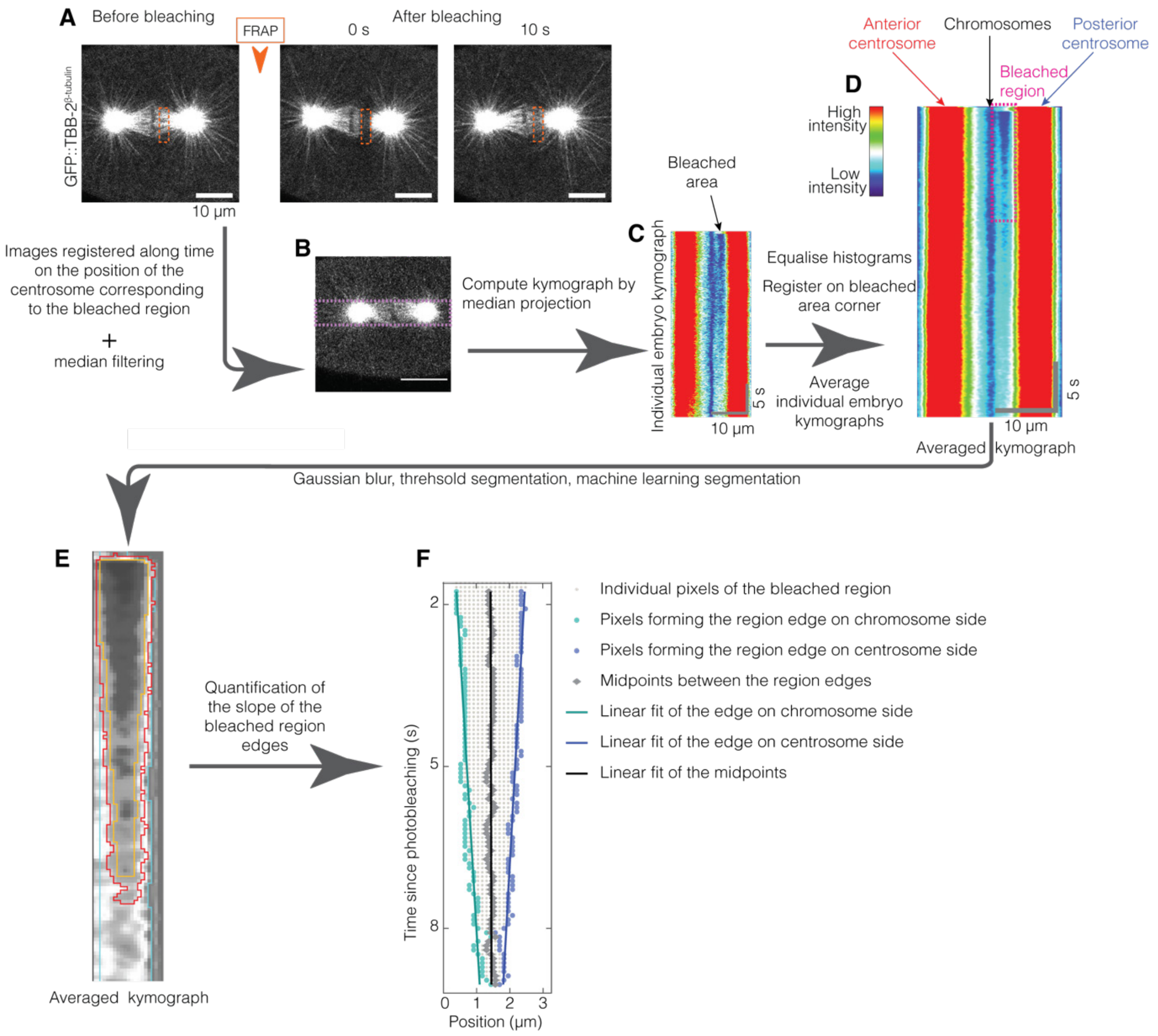


**Figure S2: Flow diagram of the image processing measuring front velocities** (**A**) Representative confocal live images before and after photobleaching. GFP::TBB-2$^{\beta\text{-tubulin}}$ labels microtubules. The scale bar corresponds to 10 µm. The orange dashed box depicts the photobleached region. (**B**) Images were registered in time on the centrosome on the same side as the bleached region, and (**C**) a kymograph was formed by median projection along the vertical dimension of the mauve dashed box superimposed to the micrograph (B). (**D**) Kymographs of the individual embryos were histogram-equalised and averaged. (**E**) Typical averaged kymograph after Gaussian blur used for segmentation. We first used a thresholding method to determine (blue contour, outside) the background pixels and (orange contour, inside) the bleached-region pixels. (red contour) We then propagated this segmentation using a random-forests machine-learning classification of the pixels (*weka*) and obtained the segmented region, post-processed using grey morphology. (**F**) This region is contoured by light blue points on the chromosome side and dark blue ones on the centrosome side, while the computed middle at each time is depicted with grey dots. We finally fitted the boundary of this region and the midline with a linear model.

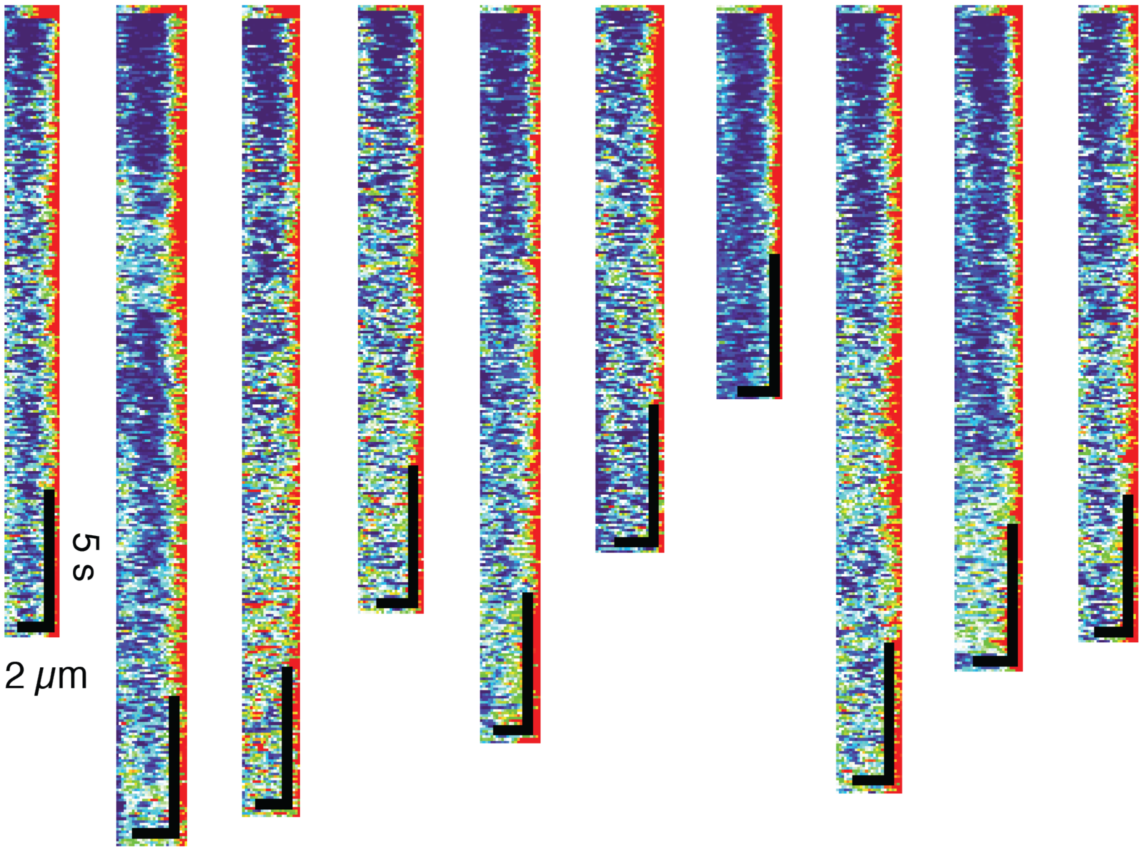


**Figure S3: Individual embryo kymographs** of the posterior half of the spindle reported for $N = 10$ GFP::TBB-2$^{\beta\text{-tubulin}}$ non-treated embryos, with centrosome on the right-hand side. They were obtained from time-registered images (Methods § Image processing). Scale bars correspond to 2 µm horizontally and 5 s vertically. These embryos are the same as in Fig 1. The colour scale ranges from blue for dark pixels to red for bright areas.

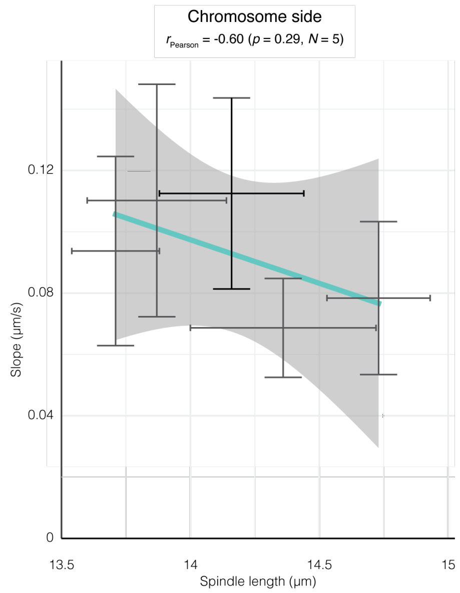


**Figure S4: Front velocity on the chromosome side did not depend on spindle length.** (**A**) The experimental front velocities on the chromosome side in (black line) non-treated and (grey line) RNAi control conditions did not show a significant correlation with the spindle length. We used strains with labelled microtubules GFP::TBB-2$^{\beta\text{-tubulin}}$. Spindle length was measured at the time of bleaching. Pearson correlation coefficient and the corresponding test are indicated above the plot. Bars correspond to means; error bars are standard errors. The grey-shaded region corresponds to the 95% confidence interval on the line coefficients.

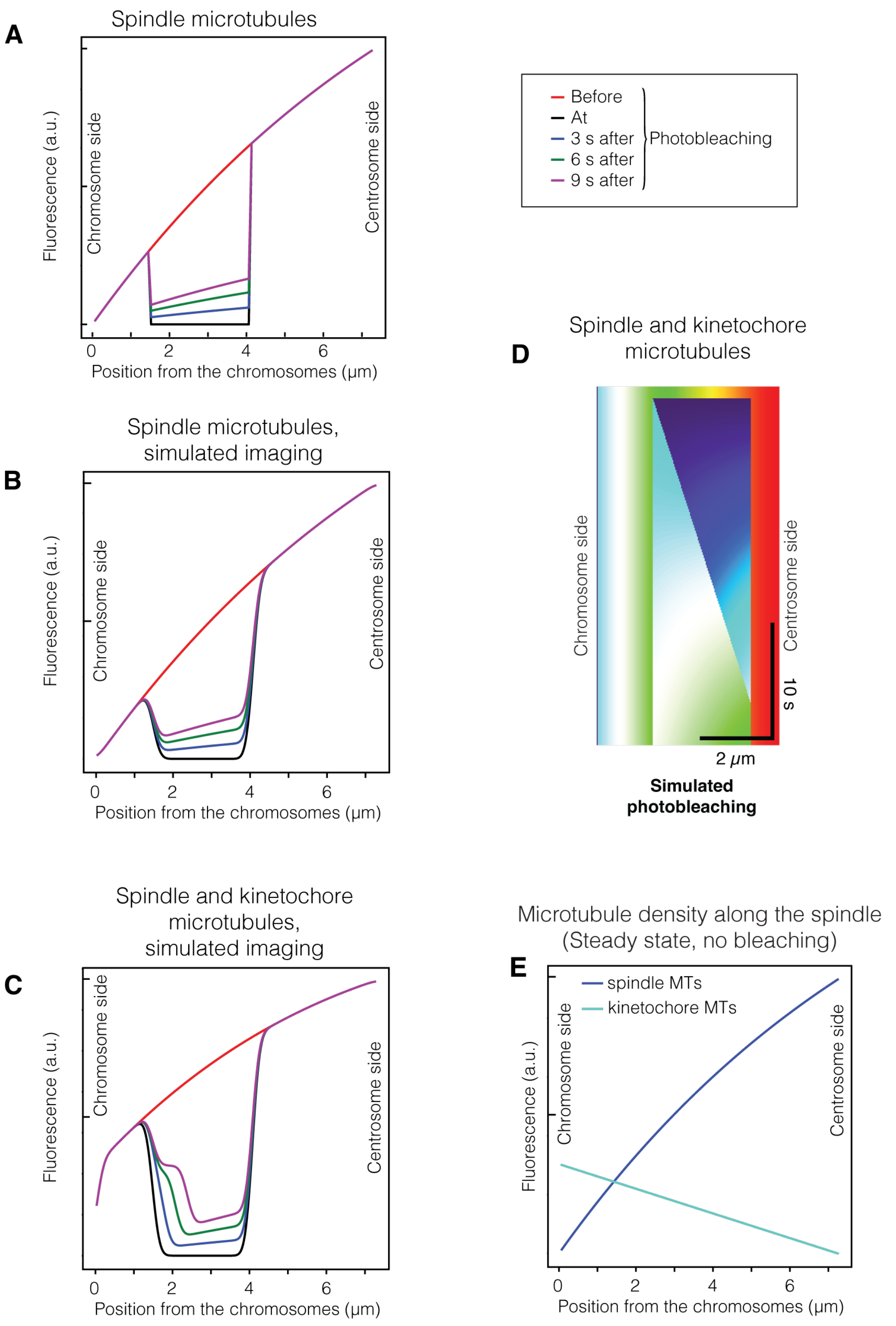


**Figure S5: Modelling the fluorescence recovery and the front displacement. (ABC)** Fluorescence profiles along the spindle (coloured lines) were modelled at various times after simulated bleaching, considering (AB) only the microtubules emanating from the centrosomes (sMTs) or (C) also the kinetochore microtubules (kMTs). The effect of microscopy imaging was (A) ignored or (BC) accounted for by convolution with the point spread function (Suppl Text §3). The portion of the curve within the bleached region was scaled by a coefficient so that the bleached fraction of tubulin underwent an exponential decay with time. (**D**) Simulated kymograph including both not-fluxing sMTs and kMTs submitted to poleward flux. We have not accounted for microscopy-imaging diffraction compared to Fig 2A done with the same simulation parameters. The colour scale ranges from blue for dark pixels to red for bright areas. (**E**) Density of sMTs and kMTs along the spindle axis, equivalent to their fluorescence in our model. Simulation parameters are reported in Suppl Table S1.

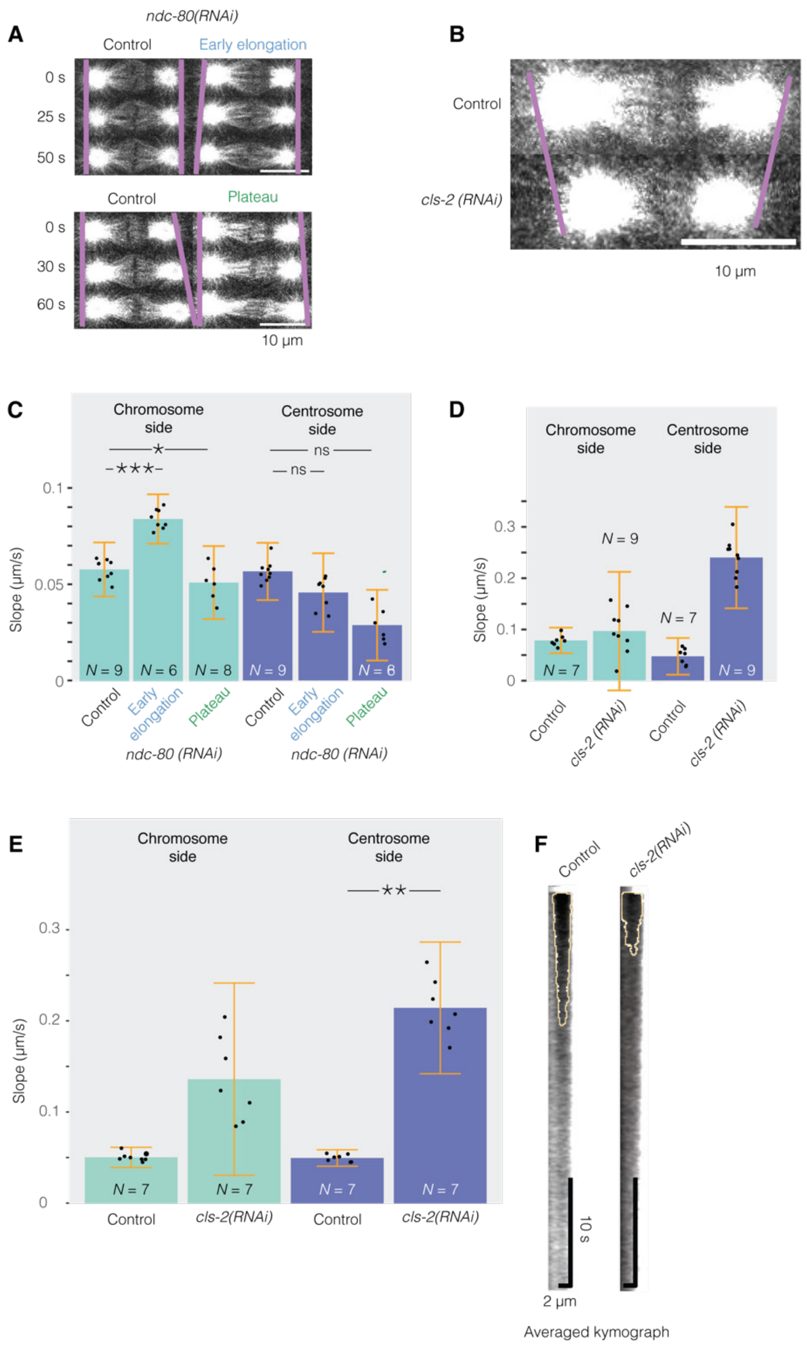


**Figure S6: Microtubule attachment and dynamics at the kinetochore impacted the front velocity. (A, B)** Exemplar micrographs of single GFP::TBB-2$^{\beta\text{-tubulin}}$ embryos used for FRAP experiment and submitted to (A) *ndc-80(RNAi),* (B) *cls-2(RNAi)* or corresponding control treatments. Times are given from the beginning of the early elongation or plateau phases. **(C, D)** Front velocities after segmenting the bleached region of: (C) $N$ = 6 *ndc-80(RNAi)* embryos bleached during precocious spindle elongation, $N$ = 8 *ndc-80(RNAi)* bleached during spindle length plateau and their $N$ = 9 controls (replica of Fig 3D); (D) $N$ = 9 *cls-2(RNAi)* treated embryos and their $N$ = 7 control ones. Microtubules were labelled using GFP::TBB-2$^{\beta\text{-tubulin}}$. Black dots represent averages of $N$-1 embryos, leaving out, in turn, each embryo (Methods § Statistics on kymograph front slopes). Bars correspond to means; error bars are estimated using Jackknife resampling. Light blue bars are values on the chromosome side and dark blue bars on the centrosome side. (**E**) Replica of the experiment (D) using $N$ = 7 *cls-2(RNAi)* treated embryos and $N$ = 7 control ones. (**F**) Averaged kymographs over the posterior half-spindle for the data presented in (D), with centrosome on the right-hand side. The orange line delineates the bleached region as obtained by our analysis (Methods § Image processing).

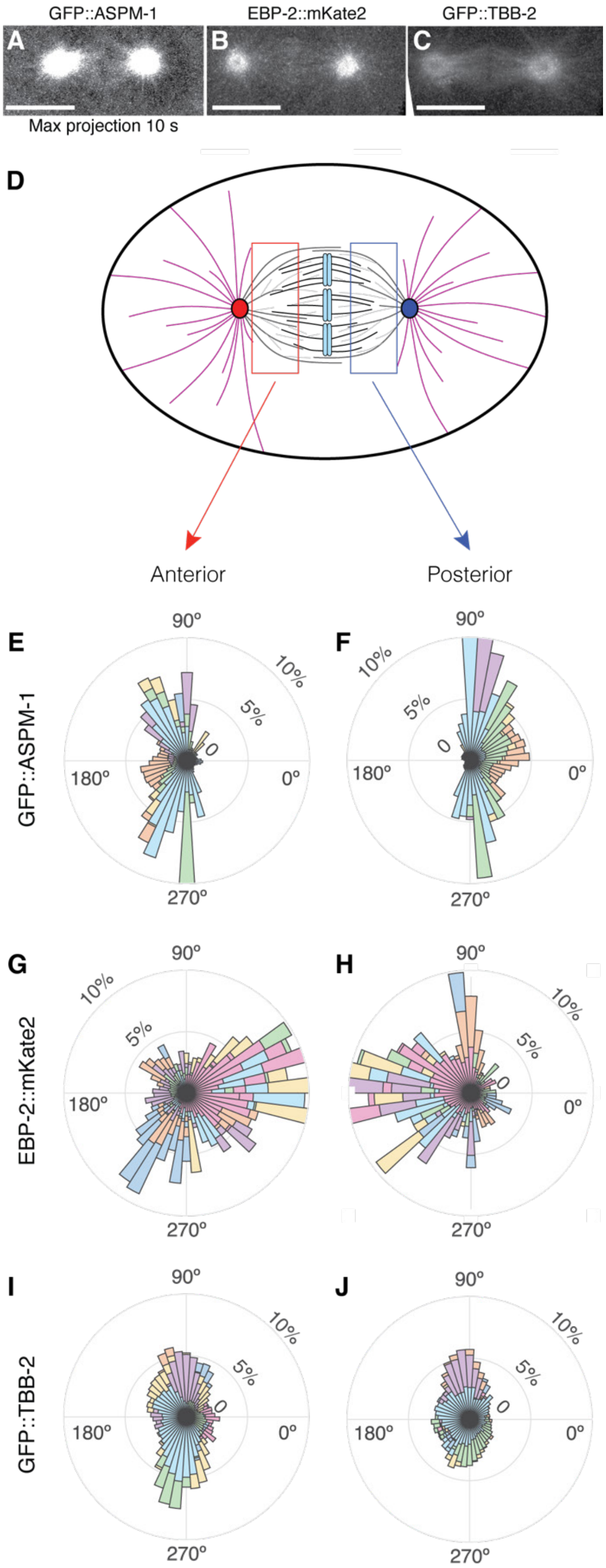


**Figure S7: Measuring the optical flow of ASPM-1::GFP, EBP-2::mKate2 and GFP::TBB-2 during metaphase.** (**A**) Typical ASPM-1::GFP embryo at metaphase, registered (Methods § Image processing), maximum projected during 10 seconds (same dataset as movie S3). The scalebar corresponds to 10 µm. (**BC**) Similar micrographs for (B) EBP-2::mKate2 and (C) GFP::TBB-2. (**D**) Schematics of the experiment to measure optical flows. Thick dark grey lines depict the spindle microtubules emanating from the (red) anterior and (blue) posterior centrosomes; thin light grey lines depict the one branching from other microtubule lattices. Black lines correspond to kinetochore microtubules bound to (blue bars) the condensed sister chromatids. Astral microtubules are depicted in purple colour. The red and blue boxes correspond to the area in which optical flow is measured in the anterior and posterior spindle halves, respectively. (**E-J**). Polar stacked histograms reporting the probability of direction of the optical flow of (EF) ASPM-1::GFP, (GH) EBP-2::mKate2, (IJ) GFP::TBB-2 comets considered in the (E, G, I) anterior and (F, H, J) posterior halves of the spindle during metaphase. Each colour corresponds to a single embryo (see Suppl. Meth.). We used $N = 6$ ASPM-1::GFP, $N = 14$ EBP-2::mKate2 and $N = 13$ GFP::TBB-2 labelled embryos. 0º corresponds to the posterior direction along the AP axis.

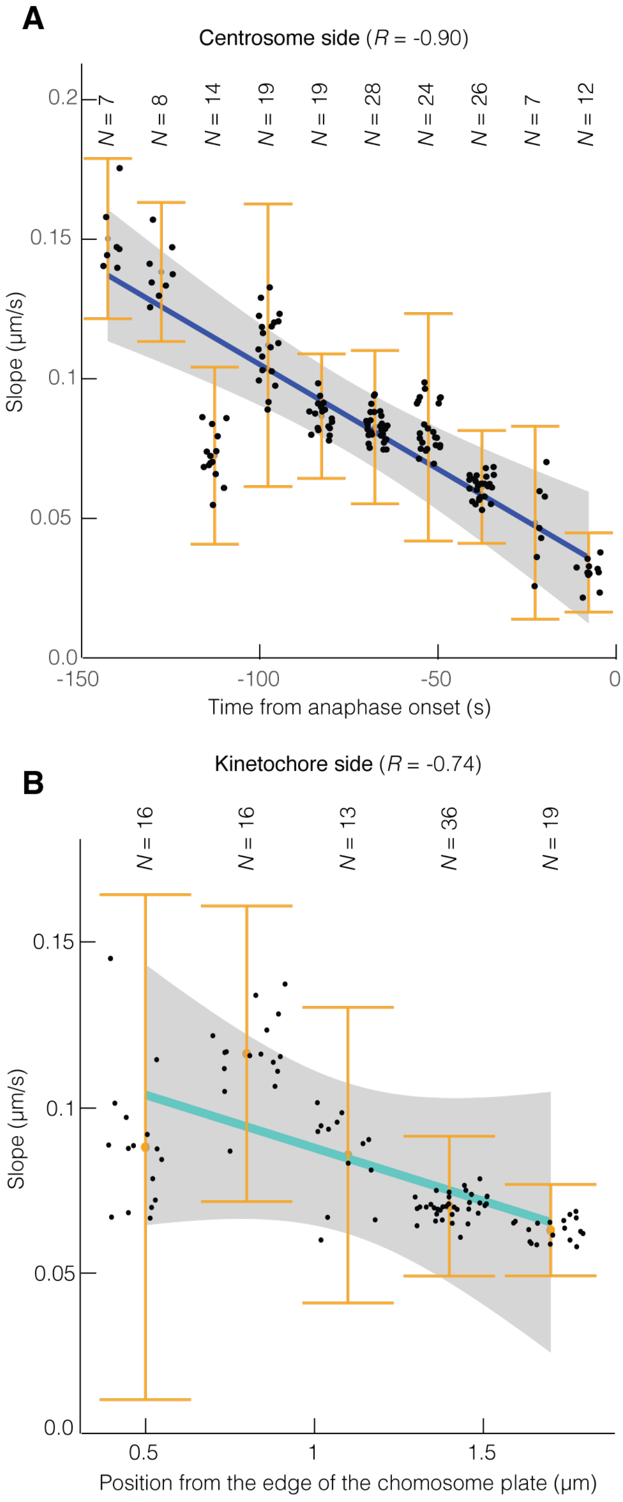


**Figure S8: Slope dependence on timing of bleaching and position along the spindle.** (**A**) Correlation between the bleaching time and the front velocities on the chromosome side. Pearson coefficient read $R = -0.90$ ($p = 0.00037$). (**B**) Correlation between the position of the chromosome-side edge of the initial bleached area and the front velocities measured on the chromosome side. Pearson coefficient read $R = -0.74$ ($p = 0.15$). In both panels, we used non-treated and control RNAi embryos of experiments reported in other figures and segmented the bleached regions. Microtubules were labelled using GFP::TBB-2$^{\beta\text{-tubulin}}$. Black dots represent averages of $N$-1 embryos, leaving out, in turn, each embryo (Methods § Statistics on kymograph front slopes). Bars correspond to means; error bars are estimated using Jackknife resampling. The grey-shaded region corresponds to the 0.95 confidence interval for the correlation-line coefficients.

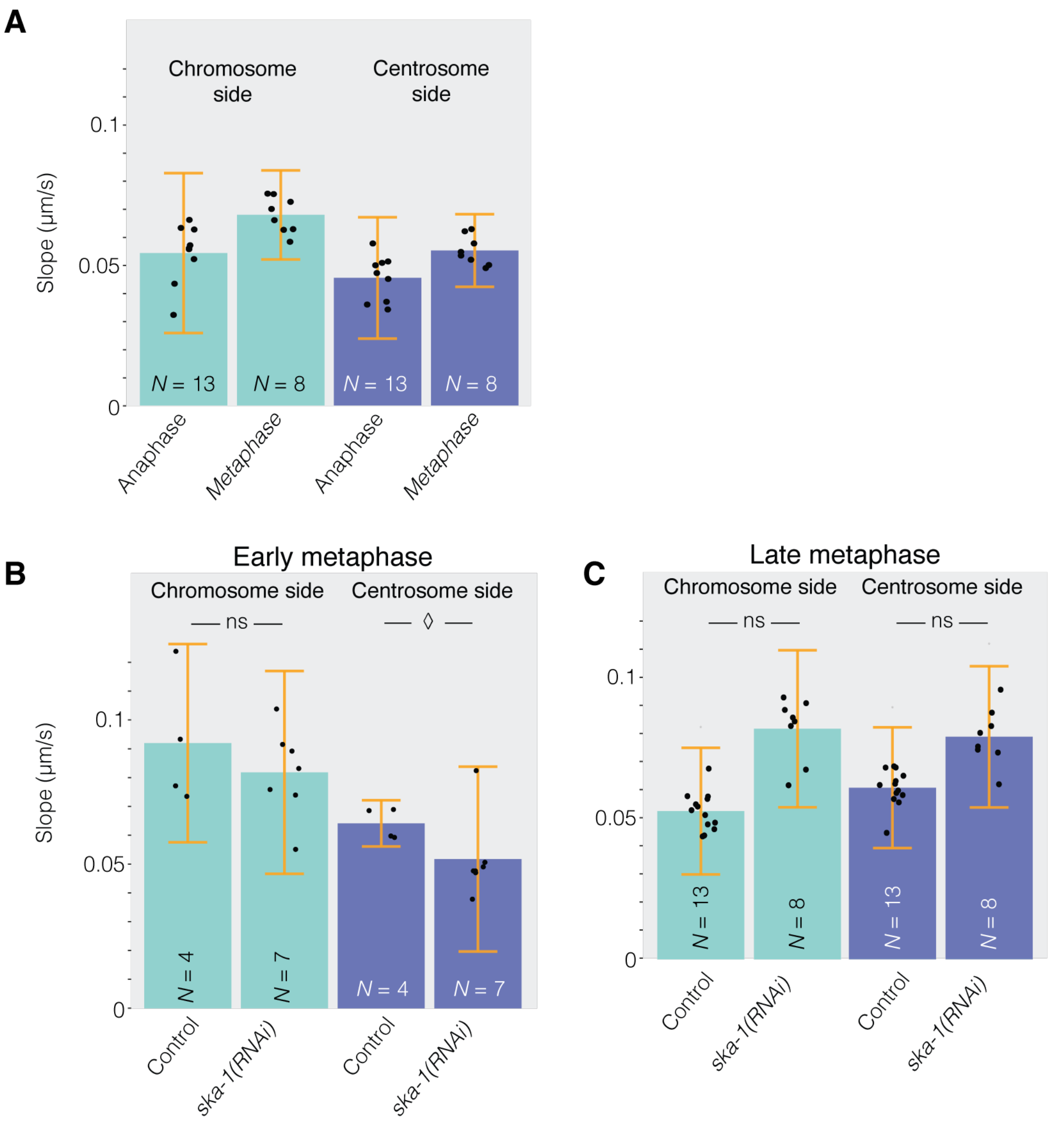


**Figure S9:** (**A**) Front velocities by segmenting the bleached region of $N = 8$ embryos bleached during metaphase and $N = 13$ during early anaphase, i.e. bleaching occurring between 10 and 20 s after anaphase onset. In all panels, microtubules were labelled using GFP::TBB-2$^{\beta\text{-tubulin}}$. (**B**) Front velocities by segmenting the bleached region of $N = 7$ *ska-1(RNAi)* embryos bleached during early metaphase (-120 to -60 s from anaphase onset) and their $N = 4$ controls. (**C**) Front velocities by segmenting the bleached region of $N = 8$ *ska-1(RNAi)* embryos bleached during late metaphase (-60 to -30 s from anaphase onset) and their $N = 13$ controls. The experiment is a replica of Fig 4B. The recovery rate was 55 ± 85% faster on the chromosome side. (BC) Black dots represent averages of $N$-1 embryos, leaving out, in turn, each embryo (Methods § Statistics on kymograph front slopes). Bars correspond to means; error bars are estimated standard errors using Jackknife resampling. Light blue bars are values on the chromosome side and dark blue on the centrosome side.

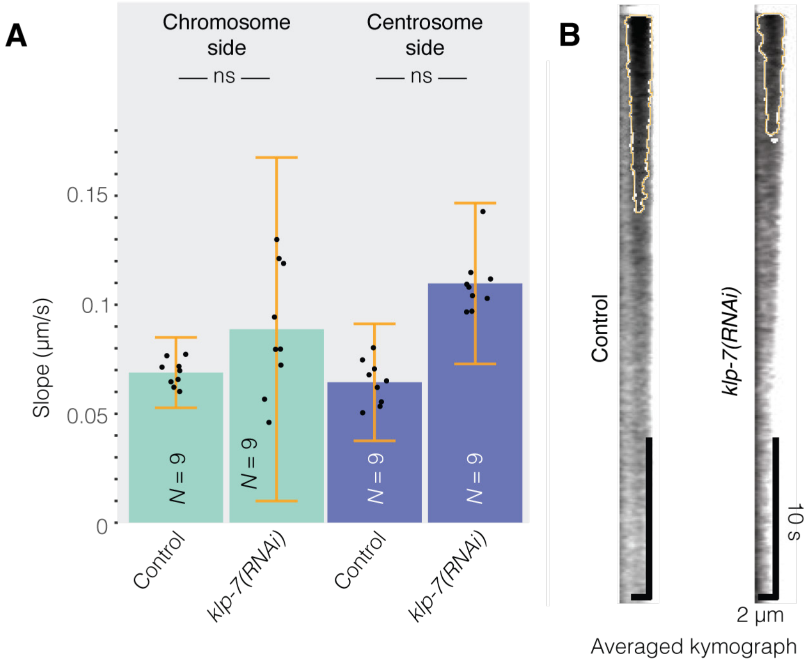


**Figure S10: KLP-7$^{MCAK}$ was dispensable for the front velocity on the chromosome side.** (A) Front velocities after segmenting the bleached region of $N = 9$ *klp-7(RNAi)* embryos and their $N = 9$ controls. Black dots represent averages of $N$-1 embryos, leaving out, in turn, each embryo (Methods § Statistics on kymograph front slopes). Bars correspond to means; error bars are estimated standard errors using Jackknife resampling. Light blue bars are values on the chromosome side, and dark blue bars correspond to the centrosome side. (B) Averaged kymograph over the posterior half-spindle for the data presented in (A), with centrosome on the right-hand side. The orange line delineates the bleached region as obtained by our analysis (Methods § Image processing). Light blue bars are velocity values measured on the chromosome side, and dark blue on the centrosome side.

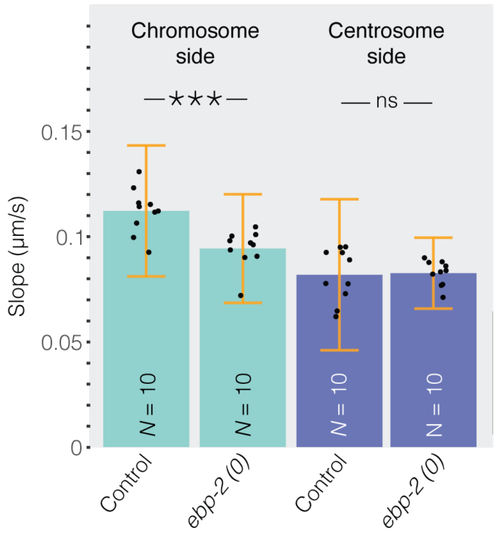


**Figure S11: EBP-2[EB1] was dispensable for the front velocity on the chromosome side.** Front velocities by segmenting the bleached region of $N$ = 10 deletion mutant *ebp-2(gk756)* embryos compared to non-treated embryos. Microtubules were labelled using GFP::TBB-2$^{\beta\text{-tubulin}}$. Black dots represent averages of $N$-1 embryos, leaving out, in turn, each embryo (Methods § Statistics on kymograph front slopes). Bars correspond to means; error bars are estimated using Jackknife resampling. Light blue bars are values on the chromosome side and dark blue bars on the centrosome side.

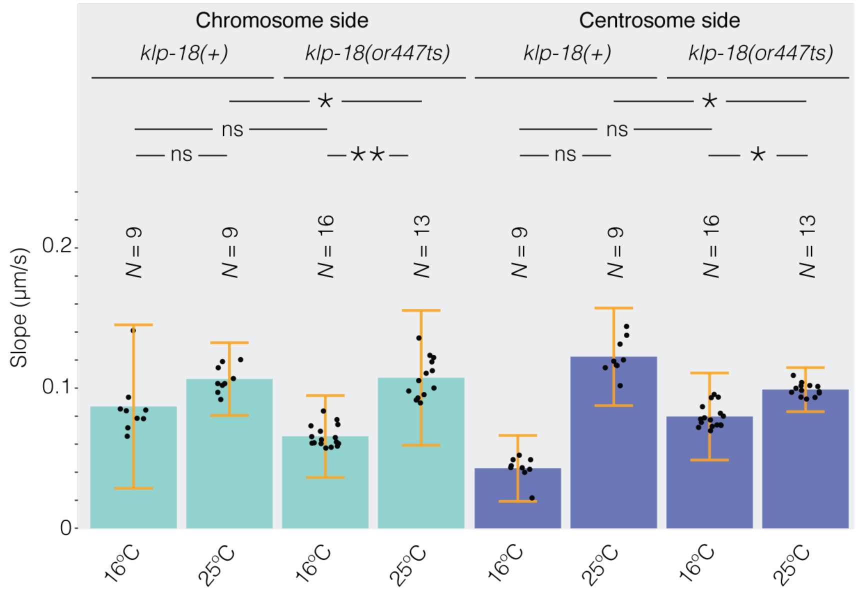


**Figure S12:** Front velocities by segmenting the bleached region of *klp-18(or447ts)* embryos at ($N$ = 16) permissive temperature 16ºC and ($N$ = 13) restrictive temperature 25ºC; corresponding control embryos klp-18(+) at ($N$ = 9) permissive temperature 16ºC and ($N$ = 9) restrictive temperature 25ºC. Microtubules were labelled using GFP::TBB-2$^{\beta\text{-tubulin}}$. The experiment was a replica of Fig 5D. Black dots represent averages of $N$-1 embryos, leaving out, in turn, each embryo (Methods § Statistics on kymograph front slopes). Bars correspond to means; errors are estimated using Jackknife resampling. Light blue bars are values on the chromosome side and dark blue on the centrosome side.

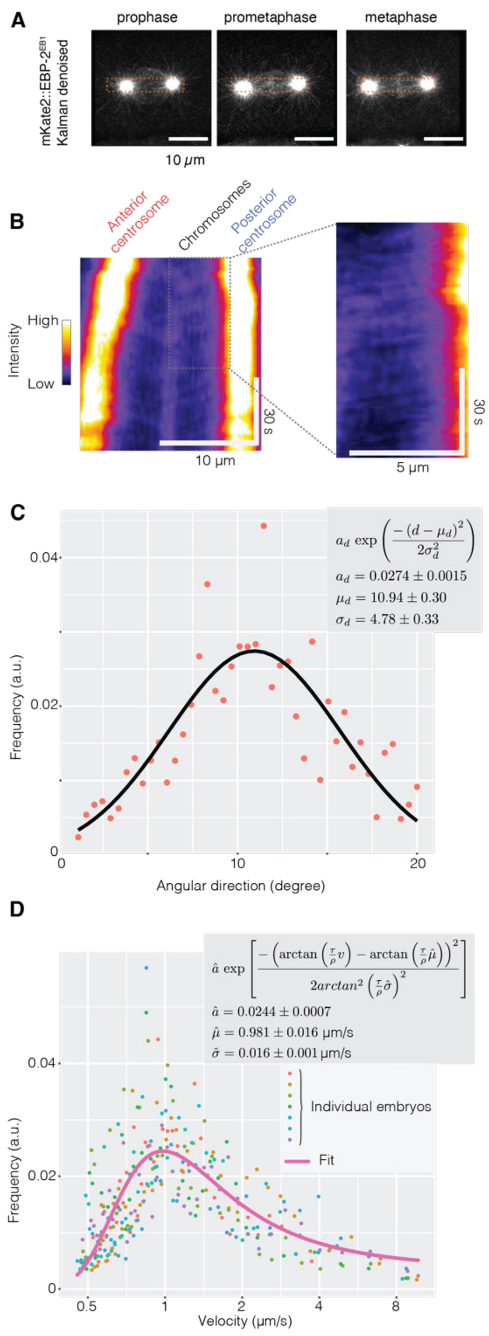


**Figure S13: Measuring microtubule growth rate in the mitotic spindle.** We imaged the spindles of $N = 7$ EBP-2$^{\text{EB1}}$::mKate2; GFP::TBB-2$^{\beta\text{-tubulin}}$ labelled embryos at 2 Hz and registered the images using the β-tubulin channel. We applied a Kalman denoising. Scale bars indicate 10 µm. (**A**) Typical stills of the EBP-2 (MT plus-ends) channel. The red dashed rectangle depicts the region used to compute the kymograph (Suppl Methods § Measuring the growth rate of microtubule plus ends). (**B**) (left part) Corresponding kymograph for a typical embryo, and (right part) magnified posterior half-spindle highlighting the traces due to microtubule growth or displacement. (**C**) (line) Exemplar fit of (dots) an individual embryo histogram of comet directions. It corresponds to the angle of the comets measured from the horizontal direction in degrees. We used the convention where positive angles correspond to counterclockwise rotation. We used a Gaussian function model, with $a$ the fit-estimated normalisation factor, $d$ the direction, $\mu_d$ the mean direction and $\sigma_d$ the corresponding standard deviation. The values obtained by fitting with standard errors are reported in the inset. (**D**) (pink line) Global fit of the comet velocity distribution of (coloured dots) individual embryos. We modelled the angular distribution of the comets with a Gaussian and transformed the direction into velocity. The equation is reported with $\hat{a}$ the fit-estimated normalisation factor, $v$ the velocity, $\tau$ the cycle time at imaging (different for each embryo), $\rho$ the pixel size (resolution), $\hat{\mu}$ the velocity and $\hat{\sigma}$ the standard deviation. The values obtained by fitting with standard errors are reported in the inset.

### Supplemental Movie

**Movie S1: Tubulin fluorescence recovery after photobleaching in the metaphasic spindle of *C. elegans*.** $N$ = 10 GFP::TBB-2$^{\beta\text{-tubulin}}$ labelled embryos were registered on the centrosome-side edge of the photobleached area and combined by computing pixel-wise median (Methods § Image processing, Fig S2). The movie plays in real-time.

**Movie S2: Localisation of KLP-18 in the spindle of a typical *C. elegans* embryo** labelled with (left) KLP-18::GFP (middle) EBP-2::mKate2 and (right) their superimposition, acquired on the spinning disk microscope (Methods § Microscopy). The movie plays in real-time.

**Movie S3: ASPM-1::GFP in the metaphasic spindle of a typical *C. elegans* embryo**, acquired on the airyscan confocal microscope with deconvolution at 18ºC and with 3.1 frames per second (Methods § Microscopy). The movie plays 2 times as fast as real time.

### Supplemental File

**File S1**: Python file computing the simulation of the front displacement due to the dynamics of the microtubules combined to the microscope diffraction (see Suppl. Text §2). It includes the iPython notebook to be displayed in JupyterLab and the YAML file to create the corresponding conda environment. Doi: 10.5281/zenodo.18772075
